# Hypothalamic oligodendrocytes regulate systemic energy balance through Notch-dependent state transitions

**DOI:** 10.64898/2026.07.29.740998

**Authors:** Simona Hankeova, Margaret Hayne, Mathijs P. Verhagen, Greg Farber, Michelle Dourado, Dewakar Sangaraju, Kayla Lee, Praveen Krishnamoorthy, Ronal Peralta, Kerstin Seidel, Kai Barck, Amy Shelton, Manal Sadek, Pamela Chan, Adel ElSohly, David Hackos, Chris Siebel, Casper Hoogenraad, Lluc Mosteiro

## Abstract

The central mechanisms through which glial cells regulate whole-body metabolism remain poorly understood. Here, we identify Notch signaling in hypothalamic oligodendrocyte lineage cells as a previously unrecognized regulator of systemic energy homeostasis. Pharmacological inhibition of the Notch ligands Jagged1 (Jag1) and Jagged2 (Jag2) induces rapid and reversible weight loss across diverse physiological and metabolic contexts independently of toxicity or caloric intake. Single-nucleus transcriptomic analyses identify hypothalamic oligodendrocyte precursor cells (OPCs) as the principal Notch-responsive population following systemic Jag1/2 inhibition and reveal expansion of a metabolically specialized GPR17⁺ intermediate state characterized by enhanced oxidative metabolism and increased predicted communication with hypothalamic neurons. This glial remodeling is accompanied by fasting-like transcriptional reprogramming of AgRP neurons, reorganization of melanocortin-autonomic circuit activity, and activation of peripheral catabolic programs. Importantly, selective deletion of Notch1/2 in hypothalamic OPCs recapitulates the major physiological and metabolic effects of systemic Jag1/2 inhibition, establishing oligodendrocyte Notch signaling as a causal regulator of whole-body metabolism. Together, our findings establish Notch-dependent oligodendrocyte state transitions as a previously unrecognized mechanism linking glial plasticity to systemic energy homeostasis.

## Introduction

Obesity and metabolic disorders pose a global health crisis, yet the central mechanisms controlling appetite and energy balance are incompletely understood^1^. The hypothalamus is a key regulator of energy homeostasis, integrating hormonal, nutrient, and neural cues to maintain body weight and metabolic balance at an organismal level^2^. Specialized neuronal populations, including Agouti-related protein (AgRP) and proopiomelanocortin (POMC) populations, orchestrate feeding behavior and energy expenditure^3–5^, while glial cells including astrocytes, oligodendrocyte precursor cells (OPCs), and tanycytes, modulate local metabolic signaling, synaptic connectivity, and neuronal activity within the hypothalamus^6–8^. While neuronal regulation of metabolism has been extensively studied, whether glial cells actively instruct systemic metabolic regulation remains largely unknown.

Notch signaling is a highly conserved pathway that regulates cell fate, intercellular communication, and tissue homeostasis across multiple organs^9,10^. Here, we show that systemic blockade of the Notch ligands Jagged1 (Jag1) and Jagged2 (Jag2) induces rapid and reversible weight loss independently of intestinal pathology, thereby uncoupling metabolic effects from the canonical goblet cell metaplasia induced by pan-Notch inhibition^11,12^. Mechanistically, Jag1/2 antibodies suppress Notch signaling in hypothalamic OPCs, resulting in expansion of a metabolically specialized GPR17⁺ intermediate state. This glial remodeling is associated with broad changes in neuronal metabolism and hypothalamic circuit activity.

## Results

### Dual Inhibition of Jag1 and Jag2 Drives Significant Weight Loss Without Intestinal Phenotypes

To investigate whether selective modulation of Notch signaling influences systemic metabolism, we examined the metabolic consequences of a unique panel of therapeutic antibodies that selectively inhibit individual Notch receptors and ligands^13–16^. We confirmed that simultaneous blockade of Notch1 and Notch2 receptors induced rapid and substantial body weight loss within one week, exceeding 20% of starting body weight (Fig. 1A). We found that dual inhibition of the Notch ligands Jag1 and Jag2 was also sufficient to induce significant weight loss (Fig. 1A). Histological analysis revealed that, as expected, dual inhibition of Notch1 and Notch2 receptors resulted in goblet cell metaplasia^17,18^ (Fig. 1B), as demonstrated by Alcian Blue staining (Extended Data Fig. 1A), accompanied by diminished Notch signaling in intestinal crypts, indicated by reduced *Hes1* expression (Fig. 1B). In contrast, blockade of Jag1 and Jag2 did not induce these intestinal phenotypic changes as both Jag ligands have limited expression in the small intestine (Fig. 1B; Extended Data Fig. 1A)^19^.

**Figure 1.**
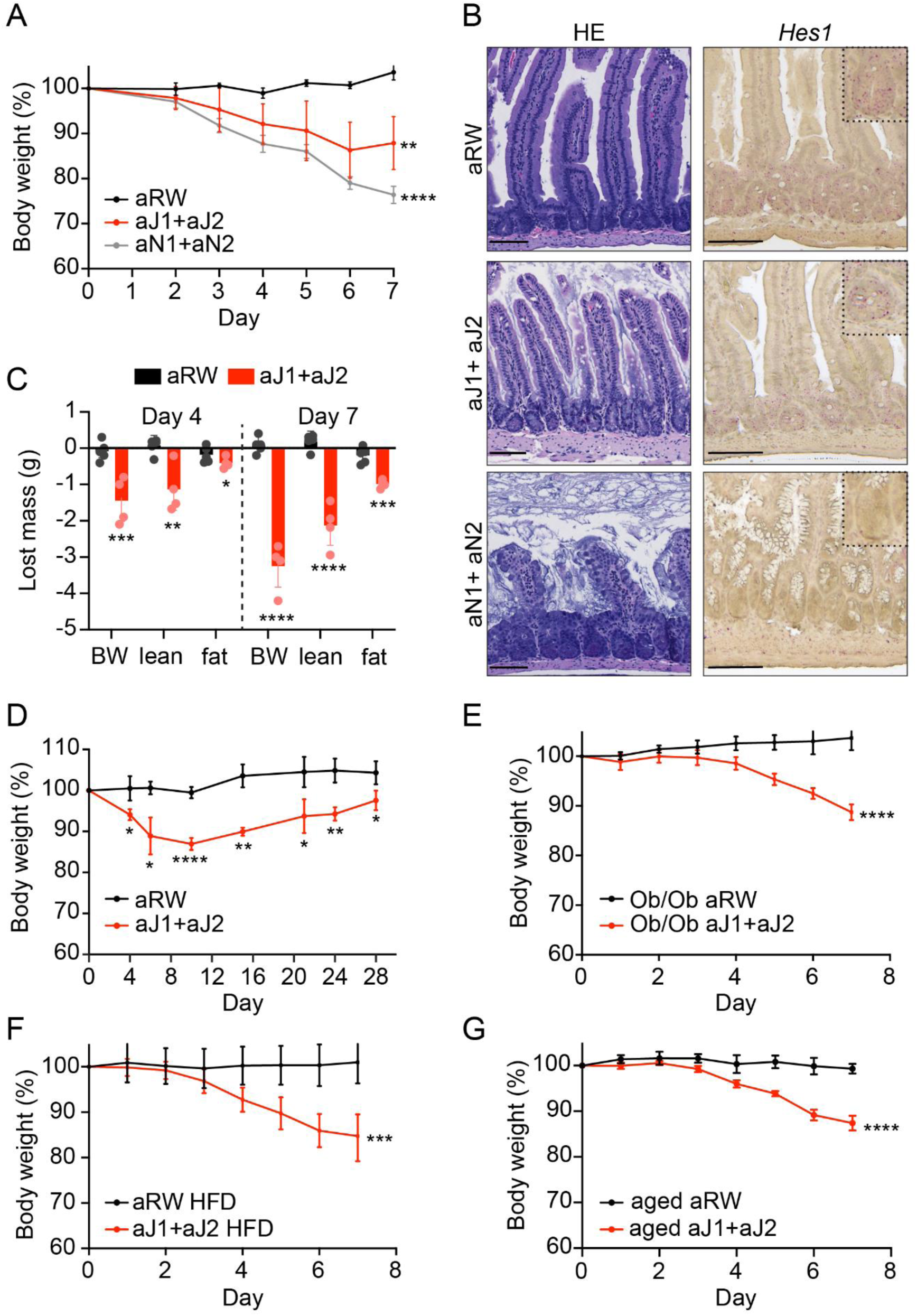
Treatment with Jag1/2 blocking antibodies causes weight loss independent of intestinal defects. A. Percentage of body weight compared to initial body weight over time up to 7 days post-treatment. Treatments were anti-Ragweed (aRW) isotype control antibody, a combination of anti-Jag1 and anti-Jag2 blocking antibodies (anti-Jag1/2), or a combination of anti-Notch1 and anti-Notch2 blocking antibodies (anti-Notch1/2). All values are expressed as mean ± s.d. of n=5 mice per group. Statistical significance was assessed by two-way ANOVA with Dunnett’s multiple comparisons test at day 7 *p*<0.01, **; *p*<0.0001, ****. B. Haematoxylin-eosin staining and RNAscope against *Hes1* in the small intestines of mice from panel A. Scale bar is 100 µm. C. Reduction in body mass (grams) compared to the initial measurement (day 0) in mice treated with anti-RW or anti-Jag1/2. The lean and fat mass was estimated from the lean and fat volumes measured by micro-computed tomography (micro-CT). All values are expressed as mean ± s.d. from at least n=4 mice per group. Statistical significance was assessed by two-way ANOVA with Sidak’s multiple comparisons test *p*<0.0, *, *p*<0.01, **; *p*<0.001, ***; *p*<0.0001, ****. BW, body weight. D. Percentage of body weight compared to initial body weight in mice after a single dose of the antibodies for 30 days. Treatments were anti-RW or anti-Jag1/2. All values are expressed as mean ± s.d. of n=5 mice per group. Statistical significance was assessed using two-way ANOVA with Sidak’s multiple comparison, *p*<0.05, *; *p*<0.01, **; *p*<0.0001, ****. E. Percentage of body weight compared to initial body weight in Ob/Ob mice across 7 days of treatment. Treatments were anti-RW or anti-Jag1/2. All values are expressed as mean ± s.d. with n=6 mice per group. Statistical significance was assessed by two-way ANOVA combined with Sidak’s multiple comparisons test at day 7 *p*<0.0001, ****. F. Percentage of body weight compared to initial body weight in high fat diet (HFD) conditions. Treatments were anti-RW or anti-Jag1/2. All values are expressed as mean ± s.d. of n=7 mice per group. Statistical significance was assessed by two-way ANOVA combined with Sidak’s multiple comparisons test at day 7 *p*<0.001, ***. G. Percentage of body weight compared to initial body weight in aged mice (> 20 months old) across 7 days. Treatments were anti-RW or anti-Jag1/2. All values are expressed as the mean ± s.d. of n=5 mice per group. Statistical significance was assessed by two-way ANOVA combined with Sidak’s multiple comparisons test at day 7 *p*<0.0001, ****.

Extensive antibody characterization has confirmed that the Jag-blocking antibodies used in these studies (anti-J1.b70 and anti-J2.b33) specifically recognize their respective ligands with negligible off-target activity^13^ (Extended Data Fig. 1B). However, the uncoupling of weight loss from intestinal phenotypes prompted development of a novel synthetic antibody, anti-J1/2.b50, which cross-reacts with both Jag1 and Jag2 and effectively inhibits signaling from these ligands (Extended Data Fig. 1B). Treatment with anti-J1/2.b50 recapitulated the rapid weight loss observed with combined anti-J1.b70 and anti-J2.b33 treatment (Extended Data Fig. 1C) and did not alter intestinal cell fate (data not shown), confirming the dissociation between weight loss and intestinal effects following Jag1/Jag2 inhibition. Importantly, treatment with a Jag1 antibody or a Jag2 antibody alone was insufficient to produce significant weight loss, suggesting a redundancy in signaling by the two ligands (Extended data Fig. 1C).

Micro-computed tomography (micro-CT) analysis revealed that weight loss reflected reductions in both lean and fat mass (Fig. 1C). Weight loss induced by combined Jag1/2 inhibition was reversible, with mice beginning to regain weight 10 days after a single administration of antibodies (Fig. 1D), which coincides with the systemic clearance of the antibodies. Furthermore, anti-Jag1/2-induced weight loss was robust across multiple physiological contexts: unaffected by genetic obesity in Ob/Ob mice^20^ (Fig. 1E) or diet-induced obesity in high-fat diet conditions (Fig. 1F), in aged mice (Fig. 1G), persisted across varying ambient temperatures (Extended Data Fig. 1D), and occurred in PPARα-deficient mice^21^ (Extended Data Fig. 1E). Notably, these changes in body weight occurred without alterations in serum cytokine levels (Extended Data Fig. 1F-1H) or peripheral blood cell counts (Extended Data Fig. 1I, 1J), ruling out systemic inflammation or hematologic toxicity.

Together, these findings identify Jag1/2-mediated Notch signaling as a regulator of systemic energy homeostasis that can be uncoupled from the intestinal toxicity associated with pan-Notch inhibition.

### Jag1/2 Inhibition Drives Coordinated Metabolic Remodeling and Energy Balance Regulation

To explore the physiological consequences of Jag1/2 blockade underlying weight loss, comprehensive metabolic phenotyping was performed. We observed that anti-Jag1/2 treatment led to coordinated reductions in food and water intake (Extended Data Fig. 2A, 2B), along with reductions in fecal and urinary output (Extended Data Fig. 2C, 2D). To determine whether reduced caloric intake accounted for the weight loss phenotype, paired-feeding experiments were performed, reducing food intake in control animals to match anti-Jag1/2-treated animals on a daily basis. Notably, only anti-J1/2-treated mice exhibited sustained weight loss despite equivalent caloric intake, indicating that reduced food consumption alone is insufficient to explain the phenotype (Fig. 2A).

**Figure 2.**
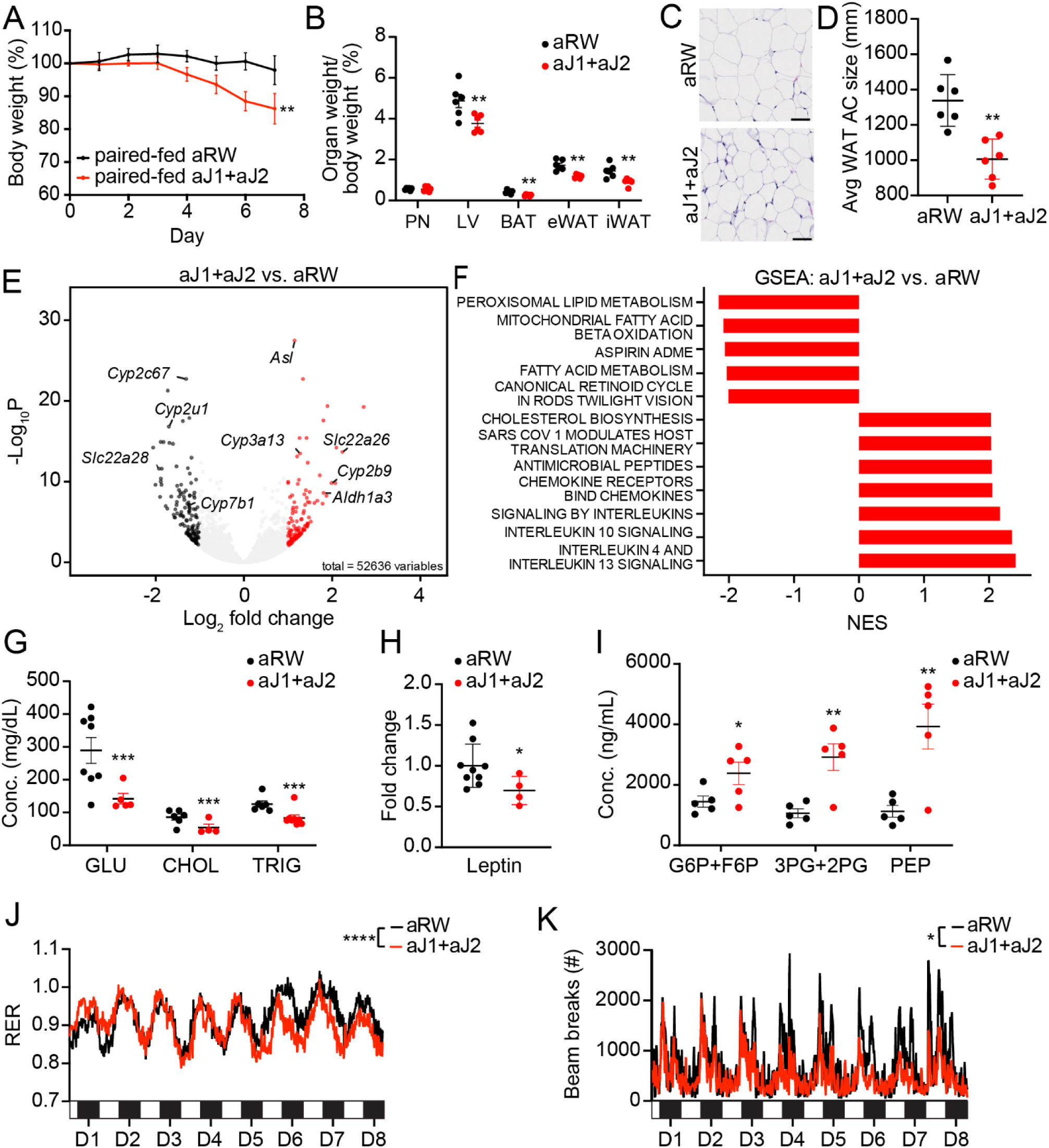
Anti-Jag1/2 treatment profoundly affects metabolism and energy homeostasis. A. Percentage of body weight compared to initial body weight in paired-fed conditions. Treatments were anti-RW or anti-Jag1/2. All values are expressed as the mean ± s.d. of n=6 mice per group. Statistical significance was assessed using two-way ANOVA with Sidak’s multiple comparison, at day 7; *p*<0.01, **. B. Percentage of wet tissue weight compared to initial body weight of paired-fed mice treated with anti-RW or anti-Jag1/2 for 7 days. All values are expressed as mean ± s.d. of at least n=5 mice per group. Statistical significance was assessed using multiple unpaired *t*-tests *p*adj<0.01, **. BAT, brown adipose tissue; eWAT, epididymal white adipose tissue; iWAT, inguinal white adipose tissue; LV, liver; PN, pancreas. C. Representative Hematoxylin-eosin images of white adipocyte tissue (WAT) of paired-fed mice treated with anti-RW or anti-Jag1/2 for 7 days. Scale bars indicate 50 μm. D. Quantification of adipocyte cell (AC) size in epididymal and inguinal white adipose tissue (WAT) sections of paired-fed mice treated with anti-RW or anti-Jag1/2 for 7 days. All values are expressed as the mean ± s.d. of n=6 mice per group. Statistical significance was assessed by Welch’s unpaired *t*-test *p*<0.01, **. E. Volcano plot showing differentially expressed genes (|log2FC| > 1.5, *p*adj < 0.01) of liver tissues of paired-fed mice treated with anti-RW or anti-Jag1/2 for 7 days of n=5 mice per group. F. Geneset enrichment analysis (GSEA) in the liver of paired-fed anti-Jag1/2-treated mice compared to anti-RW at day 7 of n=5 mice per group, using Reactome gene sets and displaying pathways with Normalized enrichment score: NES>2. G. Serum levels of glucose (Glu), cholesterol (Chol) and triglycerides (Trig) in paired-fed mice treated with anti-RW or anti-Jag1/2 for 7 days. All values are expressed as the mean ± s.d. of at least n=4 mice per group. Statistical significance was assessed by multiple unpaired *t*-test *p*adj<0.01, **; *p*adj<0.0001, ****. H. Serum leptin in paired-fed mice treated with anti-RW or anti-Jag1/2 for 7 days. All values are expressed as the mean ± s.d. of at least n=4 mice per group. Statistical significance was assessed by Welch’s *t*-test *p*<0.05, *. I. Serum levels of the indicated glycolytic intermediates in paired-fed mice treated with anti-RW or anti-Jag1/2 for 7 days. All values are expressed as the mean ± s.d. of n=5 per group. Statistical significance was assessed using multiple unpaired *t*-tests, *p*adj<0.05, *; *p*adj<0.01, **. G6P, Glucose-6-phosphate; F6P, Fructose-6-phosphate; PG, Phosphoglycerate; PEP, Phosphoenolpyruvate. J. Representation of the respiratory exchange ratio (RER) of mice treated with anti-RW or anti-Jag1/2. Each cycle represents one day and the bar represents the dark cycle (black, nighttime) and light cycle (white, daytime). All values are expressed as an average of n=7 mice per group. Statistical significance was assessed by two-way ANOVA, variation tested was treatment over time F=1.935; *p*<0.0001, ****. K. Representation of the locomotor activity of mice treated with anti-RW or anti-Jag1/2. Each cycle represents one day and the bar in the lower part represents the dark cycle (black, nighttime) and light cycle (white, daytime). All values are expressed as an average of n=7 mice per group. Statistical significance was assessed by two-way ANOVA, variation tested was treatment F=86.46, *p*<0.05, *.

Under paired-fed conditions, anti-Jag1/2 significantly reduced the weight of brown adipose tissue (BAT), consistent with prior reports of Jag1/2-mediated effects on thyroid function^22^, and was accompanied by marked reductions in white adipose tissue (WAT) and liver weights (Fig. 2B). Histological analysis of the WAT showed a decrease in the average size of adipocyte cells (Fig. 2C, 2D), consistent with increased lipolysis. The liver exhibited only minor histological changes, including sinusoidal expansion and decreased hepatocytic vacuoles, without overt morphological alterations (Extended Data Fig. 2E), and no changes in liver enzymes (Extended Data Fig. 2F). RNA sequencing of liver tissue from paired-fed mice revealed extensive transcriptional remodeling, predominantly impacting metabolic genes (Fig. 2E). Pathway analysis demonstrated enrichment in cholesterol biosynthesis (Fig. 2F), while lipid metabolic pathways were suppressed, including fatty acid oxidation and peroxisome-associated lipid metabolism, (Cyp2u1, Cyp4v3, Cyp2c67, Cyp7b1, Slc22a28) (Fig. 2E, 2F). Despite expression of Notch pathway components in the liver (Extended Data Fig. 2G), canonical Notch target genes remained unchanged (Extended Data Fig. 2H, I), suggesting that hepatic metabolic remodeling is largely indirect. Pancreatic hormones and glucose tolerance were unaltered (Extended Data Fig. 2J–M), indicating preserved pancreatic function.

Serum biochemistry and metabolomic profiling under paired-fed conditions (Extended Data Fig. 2N) revealed a systemic catabolic state characterized by reduced glucose, cholesterol, triglycerides, and leptin (Fig. 2G, H), increased circulating glycolytic intermediates (Fig. 2I), elevated β-hydroxybutyrate, and reduced isobutyrate (Extended Data Fig. 2O); while TCA intermediates and ions remained unchanged (Extended Data Fig. 2P, Q). Together, these findings indicate coordinated changes in substrate utilization characterized by enhanced lipid mobilization and altered glucose handling.

Indirect calorimetry (Oxymax) revealed a significant decrease in respiratory exchange ratio (RER) after 5 days of anti-Jag1/2 treatment (Fig. 2J), indicating a shift toward lipid utilization. This was accompanied by decreases in energy expenditure (Extended Data Fig. 2R), likely partially explained by our previously reported anti-Jag1/2-mediated thyroid dysfunction^22^, and markedly reduced spontaneous locomotor activity (Fig. 2K). Together, these findings demonstrate that Jag1/2 blockade induces a fasting-like metabolic state^23–26^ despite matched caloric intake. This uncoupling of energy balance from food intake, combined with preserved hepatic Notch signaling and pancreatic function, indicates that Jag1/2 inhibition drives systemic metabolic remodeling, likely through central mechanisms rather than direct effects in peripheral tissues.

### Jag1/2 inhibition suppresses Notch signaling in hypothalamic oligodendrocyte populations

Given the systemic metabolic effects of Jag1/2 blockade, we investigated whether the hypothalamus, a key regulator of feeding behavior and metabolic homeostasis, was a direct target of anti-Jag1/2 antibodies. Kinetic analysis of hypothalamic gene expression revealed a marked downregulation of canonical Notch targets as early as 15 h after treatment, with persistent suppression at 3 and 7 days (Fig. 3A), consistent with rapid and sustained pathway inhibition. Pharmacokinetic measurements confirmed the presence of IgG (both control and Jag1/2 antibodies) in hypothalamic tissue within the first 7 days post-injection (Fig. 3B, Extended Data Fig. 3A), which was validated by immunofluorescence detection of human IgG following systemic administration of human anti-Jag1/2 antibodies (Extended Data Fig. 3B).

**Figure 3.**
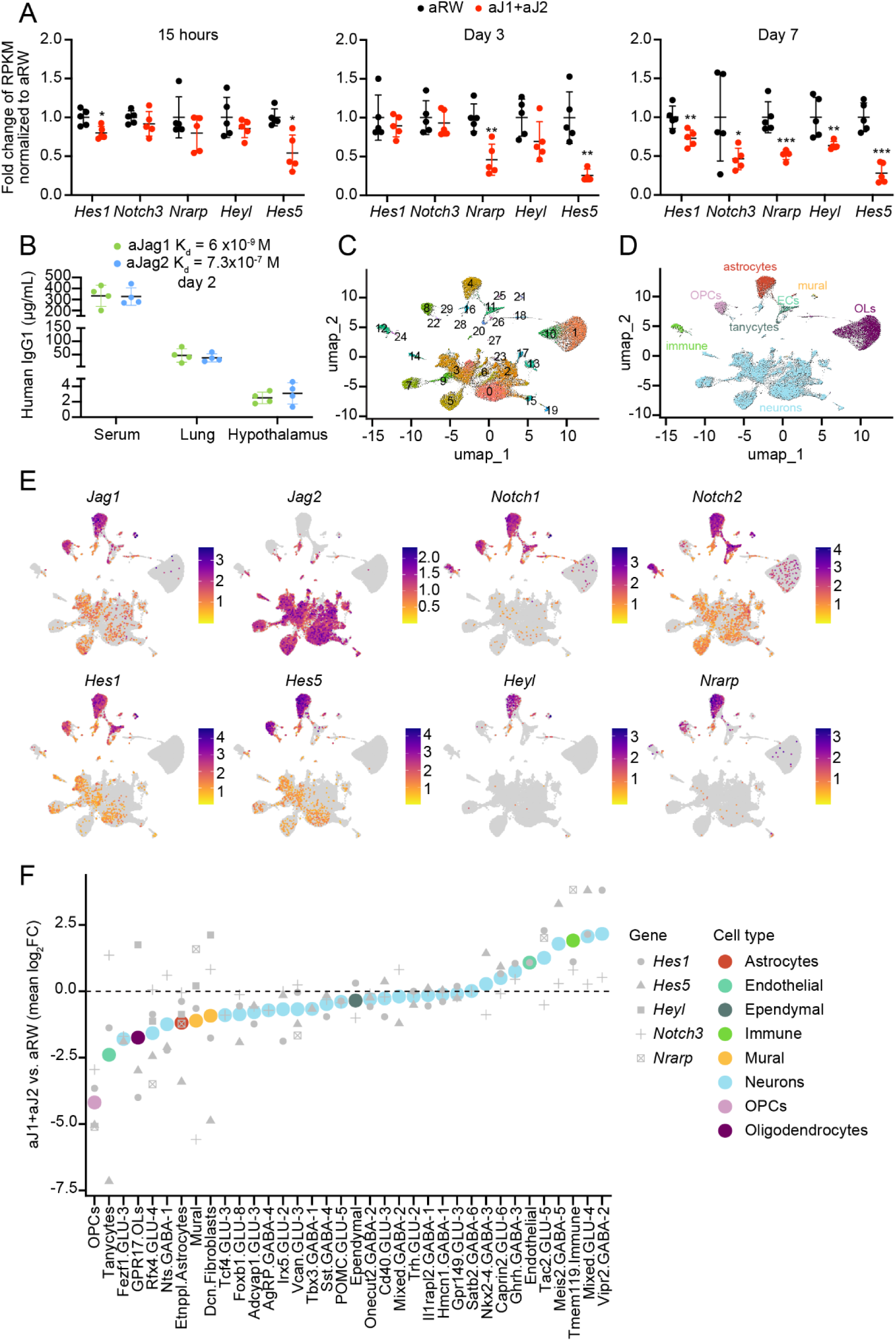
Anti-Jag1/2 downregulates Notch signaling in hypothalamic OPCs. A. Fold change expression of the indicated Notch target genes in bulk RNA-seq of murine hypothalamus treated with anti-RW or anti-Jag1/2, for 15 hours, 3 days, or 7 days. All values are expressed as mean ± s.d. of n=5 mice per group. Statistical significance was assessed by multiple unpaired *t*-test *p*adj<0.05, *; *p*adj<0.01, **; *p*adj<0.001, ***. B. Pharmacokinetic (PK) analysis of the concentration of anti-Jag1 and anti-Jag2 antibodies in the specified compartments 48 hours post-injection. All values are expressed as mean ± s.d. of n=4 mice per group. Antibody affinities (K_d_) were previously assessed by surface plasmon resonance analysis^13^. C. Uniform Manifold Approximation and Projection for dimension reduction (UMAP) plot of nuclear single-nucleus RNA-seq of the hypothalamus of ad libitum fed mice treated for 7 days with anti-RW or anti-Jag1/2, n=4 mice per group. Colors highlight the 30 clusters identified. D. Annotation of the UMAP in panel C using label transfer from the HypoMap dataset. OPCs, oligodendrocyte progenitor cells; OLs, oligodendrocytes. E. Feature plots showing scaled expression levels of Notch ligands, receptors, and target genes on the UMAP embedding. F. Downregulation of Notch target genes (*Hes1*, *Hes5*, *Hey1, Notch3* and *Nrarp*) by cell type using the C66 labels from HypoMap. Average downregulation of these genes was assessed in anti-Jag1/2-treated compared to anti-RW-treated mice. Displayed here are the mean Log_2_(FC) for Notch target genes in each cell type. OPC, oligodendrocyte progenitor cell; OLs, oligodendrocytes.

To define the cellular targets of Jag1/2 blockade, we performed single-nucleus RNA sequencing (snRNA-seq) of the hypothalamus in ad libitum fed mice treated with RW or anti-Jag1/2. UMAP embedding revealed 30 transcriptional clusters (Fig. 3C). Using HypoMap^27^ annotation, we identified cell populations corresponding to neurons, astrocytes, oligodendrocyte precursor cells (OPCs), mature oligodendrocytes (OLs), endothelial cells, vascular leptomeningeal cells, tanycytes, ependymocytes, and immune cells (Fig. 3D, Extended Data Fig. 3C - 3E). Expression mapping revealed that *Jag1* transcripts were predominantly found in astrocytes and OPCs, whereas *Jag2* expression localized primarily to neurons (Fig. 3E, Extended Data Fig. 3F, 3G). Notch receptors and target genes were mostly expressed in astrocytes and OPCs (Fig. 3E; Extended Data Fig. 3F, 3G). Strikingly, OPCs in anti-Jag1/2-treated mice exhibited the most robust downregulation of canonical Notch target genes *Hes1*, *Hes5*, *Heyl*, *Notch3* and *Nrarp* (Fig. 3F), demonstrating direct Notch pathway suppression in these cells. Other cell types showed downregulation of some, but not all Notch targets interrogated; for instance, tanycytes and fibroblasts strongly downregulated *Hes5* while mural cells downregulated *Notch3* (Fig. 3F).

These findings identify the hypothalamus as a major site of Jag1/2 antibody activity and reveal glial populations, particularly OPCs, as prominent targets. The selective suppression of Notch signaling identifies hypothalamic OPCs as the principal cellular target of Jag1/2-mediated Notch inhibition, providing the mechanistic framework linking Jag1/2 blockade to the coordinated metabolic and behavioral phenotypes characterized in Figure 2.

### Jag1/2 blockade expands metabolically specialized GPR17⁺ oligodendrocyte intermediates

To define the effects of the anti-Jag1/2 treatment in the hypothalamus independently of secondary changes in food intake, we performed single-nucleus RNA-seq analysis on a paired-fed cohort, in which control mice were food-restricted to match the reduced intake of anti-Jag1/2–treated animals. Following HypoMap-based annotation (Extended Data Fig. 4A), we confirmed that OPCs remained the cell population with the most pronounced downregulation of canonical Notch target genes (Extended Data Fig. 4B), consistent with our ad libitum dataset (Fig. 3F). We next assessed whether anti-Jag1/2 treatment altered hypothalamic cell-type abundance in either feeding condition. UMAP visualization showed broadly conserved clustering across treatment groups (Extended Data Fig. 4C) but revealed a striking expansion of GPR17⁺ oligodendrocytes in anti-Jag1/2-treated mice in both paired-fed and ad libitum conditions (Extended Data Fig. 4D, 4E). Relative abundance analysis using MiloR^28^ confirmed the pronounced enrichment of GPR17⁺ cells (Fig. 4A and Extended Data Fig. 4F). Together with suppression of canonical Notch targets in OPCs (Fig. 3F; Extended Data Fig. 4B), these findings indicate enhanced OPC-to-OL differentiation. This transition was confirmed in vivo by flow cytometry (Extended Data Fig. 4G) and *in vitro*, where anti-Notch1/2 treatment promoted OPC differentiation into OLs (Extended Data Fig. 4H), consistent with the established role of Notch in restraining OPC maturation^29–32^ while extending prior evidence that GPR17⁺ cells represent transient intermediates primed for terminal differentiation^33^.

**Figure 4.**
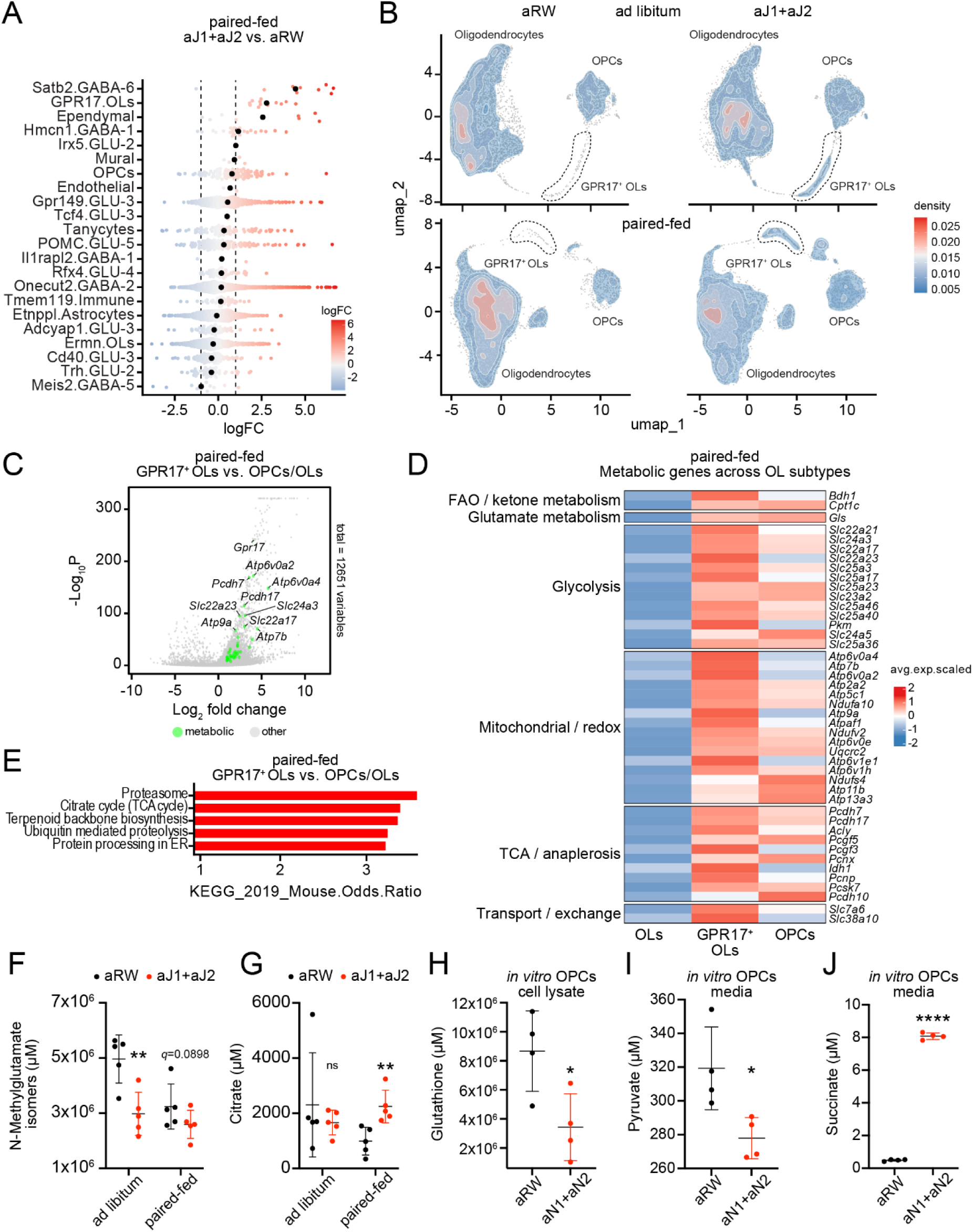
Notch downregulation induces oligodendrocyte differentiation into GPR17^+^ cells. A. Differential abundance of cells per cluster identified in snRNA-seq of the hypothalamus of paired-fed mice treated with anti-Jag1/2 compared to those treated with anti-RW for 7 days, analyzed by MiloR. Number of animals n=4 per group. OLs, oligodendrocytes: OPCs, oligodendrocyte progenitor cells. B. Density plots of cells in the UMAP of the oligodendrocyte lineage of ad libitum (top) or paired-fed (bottom) in the hypothalamus of mice treated with anti-RW or anti-Jag1/2 for 7 days. Number of animals n=4 per group. OLs, oligodendrocytes: OPCs, oligodendrocyte progenitor cells. C. Differentially expressed genes in paired-fed GPR17^+^ oligodendrocytes compared to OPCs and OLs of mice treated with anti-RW or anti-Jag1/2 for 7 days. Number of animals n=4 per group. OLs, oligodendrocytes: OPCs, oligodendrocyte progenitor cells. D. Heatmap of the differentially expressed metabolic genes in paired-fed GPR17^+^ oligodendrocytes compared to OPCs and OLs. Metabolic genes are highlighted in green. Number of animals n=4 per group. FAO, fatty acid oxidation; OLs, oligodendrocytes: OPCs, oligodendrocyte progenitor cells. E. Pathway analysis of the differentially upregulated genes in paired-fed GPR17^+^ oligodendrocytes compared to OPCs and OLs. Differentially upregulated genes with *p*adj<0.01 and log_2_FC>0.5 were selected and pathway analysis was assessed using the KEGG database. Number of animals n=4 per group. ER, endoplasmic reticulum; OLs, oligodendrocytes: OPCs, oligodendrocyte progenitor cells. F. Levels of N-Methylglutamate isomers in the hypothalamus of ad libitum and paired-fed mice treated with anti-Jag1/2 compared to those treated with anti-RW for 7 days. All values are expressed as the mean ± s.d. of n=5 mice per group. Statistical significance was assessed by Multiple unpaired *t*-test *p*adj<0.01, **. G. Levels of Citrate in the hypothalamus of ad libitum and paired-fed mice treated with anti-Jag1/2 compared to those treated with anti-RW for 7 days. All values are expressed as the mean ± s.d. of n=5 mice per group. Statistical significance was assessed by Multiple unpaired *t*-test *p*adj<0.01, **. H. Levels of Glutathione in isolated OPCs treated *in vitro* with anti-Notch1/2 or anti-RW for 3 days. All values are expressed as the mean ± s.d. of n=4 wells per group. Statistical significance was assessed by Welch’s *t*-test *p*<0.05, *. I. Levels of secreted Pyruvate in isolated OPCs treated *in vitro* with anti-Notch1/2 or anti-RW for 3 days. All values are expressed as the mean ± s.d. of n=4 wells per group. Statistical significance was assessed by Welch’s *t*-test *p*<0.05, *. ExC, extracellular. J. Levels of secreted Succinate in isolated OPCs treated *in vitro* with anti-Notch1/2 or anti-RW for 3 days. All values are expressed as the mean ± s.d. of n=4 wells per group. Statistical significance was assessed by Welch’s *t*-test *p*<0.0001, ****. ExC, extracellular.

To further define how anti-Jag1/2 treatment reshapes the OL lineage, we re-clustered OL populations. This analysis revealed substantial transcriptional heterogeneity among HypoMap-defined mature OLs (Extended Data Fig. 4I, 4J) that was not explained by differences in cell cycle progression (Extended Data Fig. 4K). Relative abundance analysis confirmed the pronounced enrichment of GPR17⁺ cells in anti-Jag1/2–treated mice (Fig. 4B; Extended Data Fig. 4J), with only modest changes in OPCs or OL subclusters (Extended Data Fig. 4J). Differential expression analysis revealed coordinated upregulation of pathways supporting substrate utilization and mitochondrial energy production, including glycolysis, oxidative phosphorylation, fatty acid and ketone metabolism, and glutamate anaplerosis (Fig. 4C–E). Together, these findings identify GPR17⁺ oligodendrocytes as a metabolically distinct intermediate state characterized by enhanced bioenergetic and anaplerotic activity. This profile supports an emerging model in which oligodendrocyte-lineage cells contribute to neuronal function through coupled glycolytic and oxidative metabolism^34–36^.

Consistent with these transcriptional changes, metabolomic profiling of the hypothalamus under both paired-fed and ad libitum conditions (Extended Data Fig. 4L) revealed reductions in N-methylglutamate isomers (Fig. 4F), whereas citrate levels were increased selectively under paired-fed conditions following anti-Jag1/2 treatment (Fig. 4G). In parallel, isolated OPCs treated with anti-Notch1/2 exhibited decreased glutathione levels (Fig. 4H), a major intracellular reservoir of glutamate, as well as reduced extracellular pyruvate (Fig. 4I) and increased extracellular succinate (Fig. 4J), consistent with increased glutamate utilization to fuel α-ketoglutarate production and TCA cycle activity. The selective accumulation of citrate under paired-feeding further suggests that this metabolic rewiring interacts with systemic nutrient availability to shape downstream carbon utilization.

Together, these findings identify GPR17⁺ oligodendrocytes as a metabolically specialized intermediate state expanded by Notch inhibition.

### GPR17⁺ oligodendrocyte expansion is associated with neuronal metabolic remodeling and **altered hypothalamic circuit activity**

Given the central role of OLs as metabolic supporters of neuronal function, we next investigated OL–neuron communication in the hypothalamus. Cell–cell interaction analysis using CellChat^37^ revealed that OPCs and GPR17⁺ oligodendrocytes represent the primary signaling-competent states within the OL lineage, exhibiting markedly higher interaction strength with multiple hypothalamic neuronal populations, including AgRP and POMC neurons (Fig. 5A, 5B), compared to mature OLs. These interactions are associated with coordinated expression of glutamate synthesis and transport machinery (*Slc1a1*, *Gls*) and engagement of diverse neuronal glutamate receptors, including AMPA, NMDA, kainate, and metabotropic receptors (Fig. 5C; Extended Data Fig. 5A). This broad receptor engagement suggests that GPR17⁺ oligodendrocytes are well positioned to modulate neuronal excitability across hypothalamic circuits and that oligodendrocyte precursor and intermediate states may preferentially engage in neuronal signaling. To identify neuronal populations most responsive to anti-Jag1/2 treatment, we performed Augur analysis^38^, which prioritizes cell types based on their transcriptional responsiveness. This analysis identified agouti-related protein (AgRP) neurons, key regulators of feeding and energy expenditure^39^, as the most responsive neuronal population in the hypothalamus (Fig. 5D; Extended Data Fig. 5B-D), indicating a preferential sensitivity of this key energy-balance circuit to the Jag1/2 blockade.

**Figure 5.**
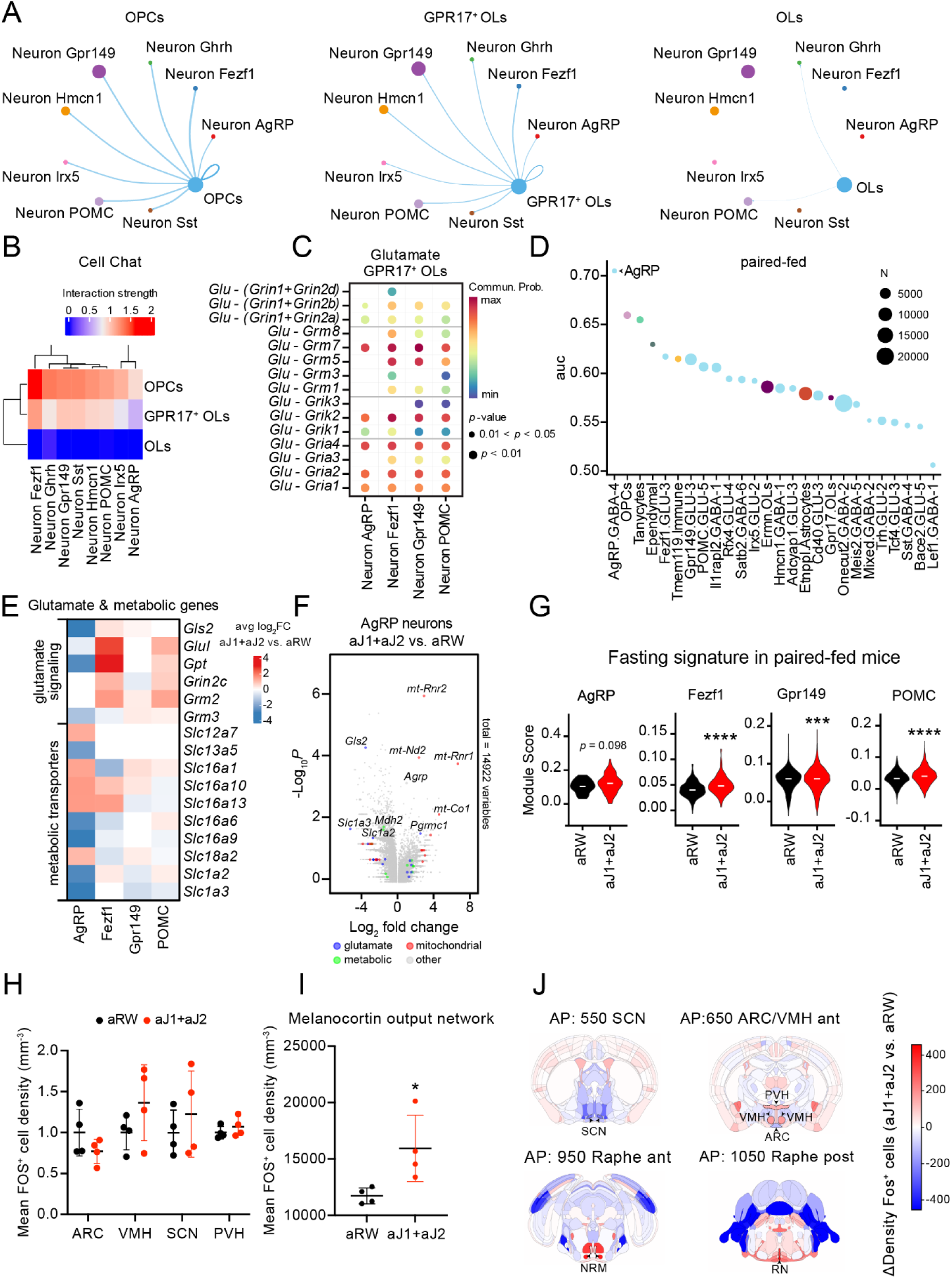
GPR17^+^ oligodendrocytes rewire the metabolism of neurons and induce a fasting-like response. A. Cell communication nodes analyzed by CellChat using the non-proteins interactome database. Circles represent the clusters and line thickness, the strength of the communication. n=4 mice per group. OLs, oligodendrocytes; OPCs, oligodendrocyte progenitor cells. B. Heatmap showing CellChat interaction strengths from OPCs, GPR17^+^ Oligodendrocytes and OLs (senders) to different neuronal populations (receivers). n=4 mice per group. OLs, oligodendrocytes; OPCs, oligodendrocyte progenitor cells. C. Dot plot of ligand-receptor communication pairs (Y axis) in the glutamate pathway identified by CellChat using the non-proteins interactome databases. Ligands are expressed in GPR17^+^ oligodendrocytes and receptors in either AgRP, Fezf1, GPR149 or POMC neurons (X axis). Circles size represents *p* value and circle color represents Communication probability. n=4 mice per group. Glu, Glu-(Slc1a1+GLS); OLs, oligodendrocytes. D. Augur analysis of the snRNA-seq data from paired-fed mice and annotated according to C66 HypoMap. Clusters with cell numbers < 20 were excluded from the analysis. Y axis represents the Area Under the Curve (AUC) of the transcriptomic changes between treatments. Dot size indicates the number of cells per cluster. Color represents annotated cell type, n=4 mice per group. OLs, oligodendrocytes; OPCs, oligodendrocyte progenitor cells. E. Heatmap of differentially deregulated genes related to glutamate metabolism and transport and transporters of key glia-neuron metabolites, in anti-Jag1/2-versus anti-RW-treated paired-fed mice in the AgRP, Fezf1, GPR149 and POMC neurons. n=4 mice per group. Color indicates average log_2_FC (red: increased; blue: decreased). F. Volcano plot of the differentially expressed genes in the AgRP neurons of anti-Jag1/2-treated paired-fed animals compared to anti-RW-treated. n=4 mice per group. Genes related to glutamate metabolism are highlighted in blue; metabolic genes in green; and mitochondrial genes in red. G. Violin plot of the fasting signature described in the HypoMap in AgRP, Fezf1, GPR149 and POMC neurons of paired-fed mice treated with anti-RW or anti-Jag1/2 for 7 days. n=4 mice per group. Statistical significance was assessed using Welch’s *t*-test, *p*<0.001, ***; *p*<0.0001, ****. H. Quantification of tdTomato+ cells density in the melanocortin-related hypothalamic nuclei of Fos-Trap (cFos-CreERT2; LSL-tdTomato) mice treated with anti-RW or anti-Jag1/2 for 4 days. n=4 mice per group. Statistical significance was assessed using Mann-Whitney test. I. Quantification of tdTomato+ cells density in the melanocortin output nuclei of Fos-Trap (cFos-CreERT2; LSL-tdTomato) ad libitum fed mice treated with anti-RW or anti-Jag1/2 for 4 days. The graph represents the sum of melanocortin output hypothalamic nuclei: paraventricular (PVH) together with melanocortin output brainstem nuclei (parabrachial, nucleus of the solitary tract, dorsal motor nucleus of the vagus nerve, medulla and nucleus Raphe). n=4 mice per group. Statistical significance was assessed using Mann-Whitney test. *p*<0.05, *. J. Brain maps of the indicated brain areas showing the tdTomato+ cells density in anti-Jag1/2 compared to anti-RW mice, treated for 4 days. Red indicated increased tdTomato+ cells density and blue, decreased density. Ant, anterior; ARC, arcuate nucleus; NRM, nucleus raphe magnus; post, posterior; PVH, paraventricular hypothalamic nucleus; RN, nucleus raphe magnus and pallidus; SCN, suprachiasmatic nucleus; VMH, ventromedial nucleus of the hypothalamus.

We next examined transcriptional changes across the top four Augur-ranked neuronal subtypes (AgRP, Fezf1, GPR149, and POMC neurons) and observed broad remodeling of metabolic programs following anti-Jag1/2 treatment, with the strongest effects in AgRP neurons (Fig. 5E, 5F; Extended Data Fig. 5E–I). Differential expression was preferentially enriched in pathways related to cellular metabolism rather than reflecting global transcriptional shifts. These changes prominently involved glutamate handling, metabolite transport, and oxidative metabolism (Fig. 5E; Extended Data Fig. 5F). In parallel, anti-Jag1/2 treatment induced coordinated upregulation of mitochondrial and bioenergetic programs across all four neuronal subtypes (Fig. 5F; Extended Data Fig. 5E–I), together with remodeling of pathways related to glucose, glutamate, fatty acid, and ketone utilization, consistent with enhanced oxidative metabolism and altered glia–neuron metabolic coupling. In line with these changes, neuronal subtypes also showed increased enrichment of the HypoMap fasting signature following anti-Jag1/2 treatment (Fig. 5G), indicating induction of a fasting-like transcriptional state independent of caloric intake. Together, these findings demonstrate that Jag1/2 blockade remodels glia–neuron metabolic interactions and is associated with increased mitochondrial activity and bioenergetic demand across multiple neuronal subtypes, with the strongest effects observed in AgRP neurons.

To functionally assess neuronal activation, we treated Fos-TRAP^40^ (cFos-CreERT2; LSL-tdTomato) mice with anti-Jag1/2 antibodies and administered 4-hydroxytamoxifen after 4 days to permanently label recently active (Fos⁺) neurons with tdTomato. We then performed 3D light-sheet fluorescence microscopy on cleared mouse brains. Quantitative imaging of hypothalamic (Fig. 5H; Extended Data Fig. 5J) and brainstem regions (Extended Data Fig. 5K) revealed a rewiring of melanocortin circuit activity, characterized by reduced neuronal activity in the arcuate nucleus (ARC; sensory node) (Fig. 5H, 5J), coupled with increased activation of downstream output regions (Fig. 5H, 5I), including paraventricular (PVH), ventromedial hypothalamus (VMH) and brainstem nuclei (parabrachial, nucleus of the solitary tract, dorsal motor nucleus of the vagus nerve, medulla and nucleus Raphe) (Fig. 5I, 5J; Extended Data Fig. 5K), consistent with enhanced autonomic output.

Together, these findings identify GPR17⁺ oligodendrocytes as the intermediary state linking Notch inhibition to neuronal metabolic remodeling and hypothalamic circuit reorganization.

### Hypothalamic OPC-specific Notch1/2 deletion establishes a causal role for oligodendrocyte Notch signaling in systemic energy homeostasis

Building on our findings that anti-Jag1/2 treatment suppresses Notch signaling in hypothalamic OPCs, expands GPR17⁺ intermediates, and remodels neuronal metabolic programs and circuit activity, we next asked whether selective loss of Notch signaling in OPCs is sufficient to recapitulate the systemic metabolic phenotypes observed with anti-Jag1/2 treatment. To this end, we stereotaxically injected AAVs encapsulating PDGFRα-Cre recombinase into the mediobasal hypothalamus of Notch1^fl/fl^; Notch2^fl/fl^ mice (Fig. 6A), thereby restricting recombination and genetic deletion of Notch1 and Notch2 in hypothalamic OPCs. Littermate controls received AAVs expressing Blue Fluorescent Protein (BFP) only. Efficient targeting was confirmed by RNAscope for BFP together with the oligodendrocyte marker Olig1, while genetic recombination was validated by reduced expression of the Notch target gene Hes5 in Olig1⁺ cells (Extended Data Fig. 6A).

**Figure 6.**
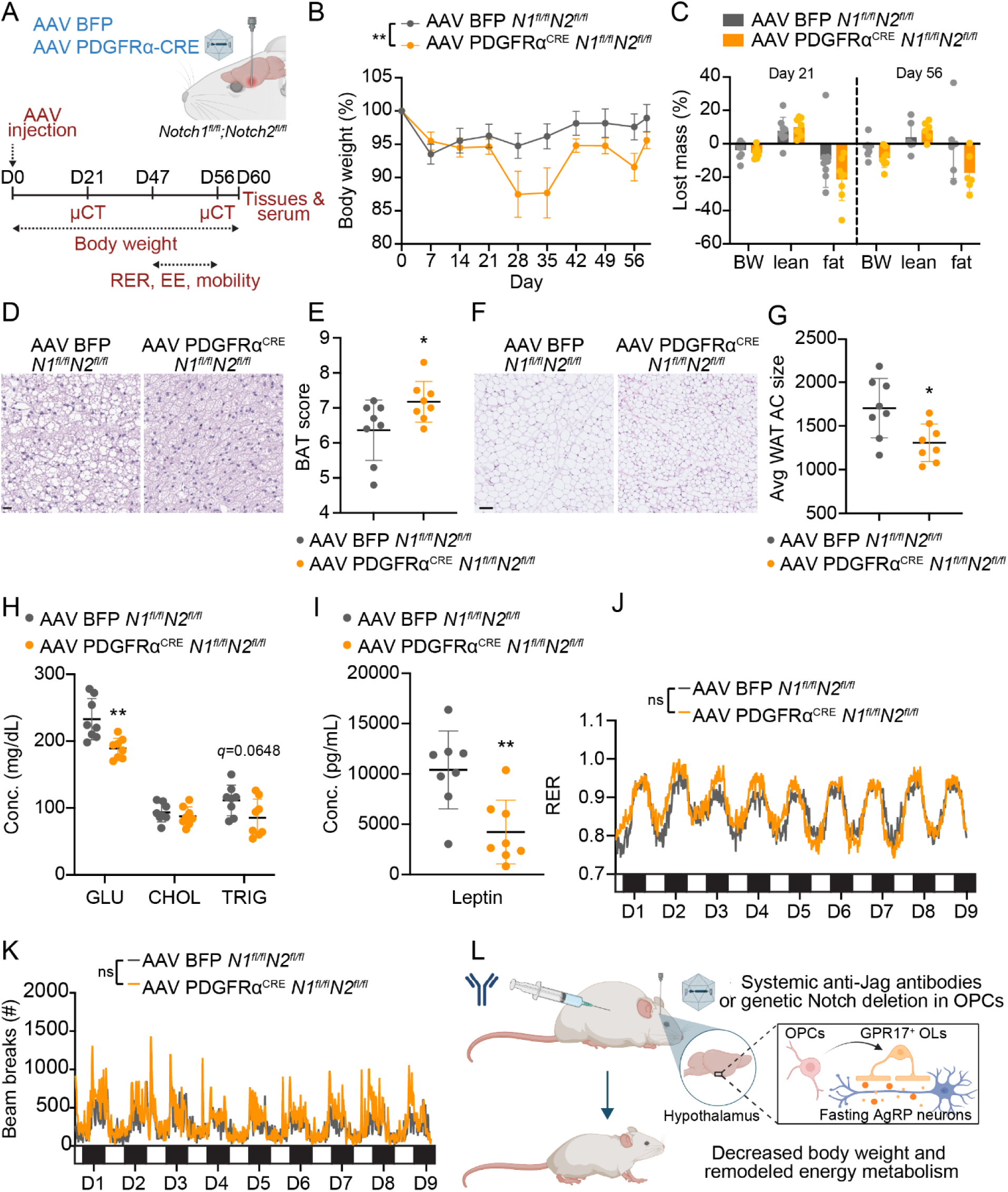
OPC-specific Notch1/2 deletion in the hypothalamus induces metabolic phenotypes similar to anti-Jag1/2 treatment. A. Schematic of the AAV stereotaxic injection and experimental set up. Designed in Biorender. B. Percentage of body weight compared to initial body weight in *Notch1*^fl/fl^; *Notch2*^fl/fl^ mice that have been stereotaxically injected with AAVs encapsulating the fluorescent reporter BFP only (CRE-) or Pdgfrα-Cre (CRE+). All values are expressed as mean ± s.d. with n=8 mice per group. Statistical significance was assessed by two-way ANOVA (time vs. genotype, *p*<0.01, **) with Sidak’s multiple comparisons test, day 35 *p*<0.05, *. C. Reduction in body mass (%) measured by micro-computed tomography (micro-CT) compared to the initial measurement (day 0) in mice from (A), measured 21 or 56 days after stereotaxic injection. All values are expressed as mean ± s.d. from at least n=6 mice per group. Statistical significance was assessed by multiple unpaired *t*-tests. D. Brown adipose tissue (BAT) activity score in mice from (A), assessed blindly. Score of 1 is white adipose tissue phenotype and score of 10 is BAT at its maximum capacity. All values are expressed as the mean ± s.d. of n=8 mice per group. Statistical significance was assessed by Welch’s *t*-test *p*<0.05, *. E. Representative Hematoxylin-eosin images of BAT of mice in (A). Scale bars indicate 20 μm. F. Quantification of adipocyte cell (AC) size in epididymal and inguinal white adipose tissue (WAT) sections of mice from (A). All values are expressed as the mean ± s.d. of n=8 mice per group. Statistical significance was assessed by Welch’s *t*-test *p*<0.05, *. G. Representative Hematoxylin-eosin images of WAT of mice in (A). Scale bars indicate 100 μm. H. Serum levels of glucose (Glu), cholesterol (Chol) and triglycerides (Trig) in mice of (A) assessed at day 59. All values are expressed as the mean ± s.d. of at least n=8 mice per group. Statistical significance was assessed by multiple unpaired *t*-test *p*adj<0.01, **. I. Serum levels of leptin in mice from (A) assessed at day 59. All values are expressed as the mean ± s.d. of n=8 mice per group. Statistical significance was assessed by Welch’s *t*-test *p*<0.01, **. J. Representation of the respiratory exchange ratio (RER) of mice from (A) during days 47-56 after stereotaxic injection. Each cycle represents one day and the bar in the lower part represents the dark cycle (black, nighttime) and light cycle (white, daytime). All values are expressed as an average of n=8 mice per group. Statistical significance was assessed by Mixed effects analysis. K. Representation of the mobility (assessed by number of beam breaks) of mice from (A) during days 47-56 after stereotaxic injection. Each cycle represents one day and the bar in the lower part represents the dark cycle (black, nighttime) and light cycle (white, daytime). All values are expressed as an average of n=8 mice per group. Statistical significance was assessed by Two-way ANOVA analysis. L. Proposed model of the effects of anti-Notch antibodies or Notch genetic deletion in the hypothalamus. Notch is suppressed in OPCs, promoting expansion of GPR17⁺ oligodendrocyte intermediates, remodeling neuronal metabolic programs, particularly AgRP neurons, and ultimately inducing a systemic catabolism state, characterized by body weight loss and remodeled energy metabolism. Designed in Biorender.

Deletion of Notch1/2 in OPCs resulted in a progressive reduction in body weight, reaching ∼5–10% below controls from day 21 through day 60 after injection (Fig. 6B), without significant changes in individual tissue weights (Extended Data Fig. 6B). Micro-CT analysis revealed a selective decrease in fat mass accompanied by a modest increase in lean mass (Fig. 6C), indicating reduced adiposity rather than generalized tissue loss. Consistent with these changes, PDGFRα-Cre mice exhibited activation of brown adipose tissue (BAT), characterized by reduced lipid droplet content and smaller adipocyte size (Fig. 6D, 6E), suggesting that the effects of systemic anti-Jag1/2 in BAT are driven, at least in part, by central mechanisms in addition to previously described anti-Jag1/2 driven thyroid defects^22^. Similarly, white adipose tissue (WAT) showed increased lipid mobilization, including reduced adipocyte size and lipid accumulation (Fig. 6F, 6G), phenocopying the effects of systemic anti-Jag1/2 in WAT (Fig. 2C, 2D). Consistent with this systemic catabolic phenotype, serum profiling revealed reduced glucose, a trend toward decreased triglycerides (Fig. 6H) and markedly decreased leptin (Fig. 6I), closely phenocopying the effects of anti-Jag1/2 treatment (Fig. 2G, 2H). In contrast, circulating cholesterol remained unchanged (Fig. 6H), and no alterations were observed in liver enzymes or serum ions (Extended Data Fig. 6C, 6D), indicating a lack of systemic toxicity. Together, these findings demonstrate activation of a systemic catabolic program downstream of hypothalamic OPC-specific Notch deletion, linking central oligodendrocyte signaling to peripheral metabolic regulation via increased activation of melanocortin-associated autonomic output circuits, as indicated by elevated Fos labeling in these regions (Fig. 5I).

Importantly, these metabolic changes occurred in the absence of differences in food intake between Cre⁺ and Cre⁻ control groups (Extended Data Fig. 6E), demonstrating that OPC-specific Notch loss regulates energy balance independently of caloric consumption, consistent with our paired-fed anti-Jag1/2 experiments (Fig. 2A). Indirect calorimetry revealed an increase in respiratory exchange ratio (RER) (Fig. 6J), particularly during the dark (active) period (Extended Data Fig. 6F), indicating altered substrate utilization and increased metabolic flexibility rather than a simple shift toward carbohydrate usage. This was accompanied by increased locomotor activity (Fig. 6K; Extended Data Fig. 6G). Energy expenditure normalized to body weight remained unchanged (Extended Data Fig. 6H, 6I), while total absolute energy expenditure was elevated in Cre-injected mice (Extended Data Fig. 6J), suggesting that the lean phenotype is associated with increased locomotor activity and altered fuel utilization rather than a primary increase in basal metabolic rate.

These findings demonstrate that hypothalamic OPC-specific deletion of Notch1/2 is sufficient to recapitulate key aspects of the metabolic phenotype induced by systemic Jag1/2 blockade, including reduced body weight, activation of thermogenic and lipolytic pathways, lowered serum glucose and leptin, and altered substrate utilization and locomotor activity. Importantly, these effects occur independently of food intake and in the absence of systemic toxicity. Collectively, these findings establish hypothalamic OPC-intrinsic Notch signaling as a causal regulator of systemic energy homeostasis and demonstrate that Notch-dependent oligodendrocyte state transitions remodel hypothalamic neuronal circuits to coordinate whole-body metabolism (Fig. 6L).

## Discussion

Obesity and metabolic disease remain major global health challenges, yet current therapeutic strategies are largely centered on peripheral metabolic tissues or neuronal appetite pathways. Although recent advances, particularly GLP-1 receptor agonists, have demonstrated the therapeutic potential of central metabolic regulation^41^, most approaches continue to focus predominantly on neuronal circuits controlling appetite and satiety. Here, we identify hypothalamic oligodendrocyte-lineage cells as active regulators of systemic energy balance, revealing a previously underappreciated glia-to-neuron axis controlling organismal metabolism.

While astrocytes and microglia have been implicated in hypothalamic nutrient sensing and inflammatory remodeling ^42,43^, oligodendrocyte-lineage cells have largely been viewed as structural or myelinating cells. Our findings establish oligodendrocyte-lineage cells as active regulators of hypothalamic metabolic function, expanding the repertoire of glial mechanisms controlling systemic energy homeostasis.

Mechanistically, we identify Jag1/2-dependent Notch signaling as a pathway maintaining hypothalamic OPC state and restraining oligodendrocyte-lineage remodeling. Jag1/2 inhibition promotes expansion of a GPR17⁺ oligodendrocyte intermediate state associated with enhanced glycolytic, mitochondrial, glutamatergic, and lipid metabolic programs. Although GPR17⁺ oligodendrocytes have primarily been studied in developmental transitions and remyelination, our findings suggest they may additionally function as metabolically specialized signaling states within adult hypothalamic circuits.

Remodeling of the oligodendrocyte lineage was accompanied by coordinated changes in neuronal metabolic programs and hypothalamic circuit activity despite minimal evidence of direct neuronal Notch inhibition, indicating that glial state transitions indirectly reshape neuronal function through metabolic coupling.

At the organismal level, oligodendrocyte-lineage remodeling activates melanocortin-associated autonomic circuits, promotes adipose tissue catabolism, and drives systemic metabolic remodeling. These observations position oligodendrocyte-lineage cells as regulators of canonical neuronal energy-balance pathways and highlight glial remodeling as an important determinant of organismal metabolism. Although systemic Jag1/2 blockade and OPC-specific Notch deletion share core metabolic phenotypes, including reduced adiposity and altered substrate utilization, differences in locomotor activity and energy expenditure likely reflect additional peripheral contributions of systemic ligand inhibition, including previously described effects on thyroid physiology^22^.

Our findings also highlight the importance of pathway-selective modulation within the Notch signaling system. Pan-Notch inhibition has long been limited by intestinal toxicity driven by disruption of epithelial differentiation^13,17,18^. In contrast, Jag1/2 blockade uncouples systemic metabolic regulation from canonical intestinal goblet cell metaplasia, demonstrating that selective ligand targeting can preserve metabolic effects while avoiding major gastrointestinal toxicity. This distinction may have broader implications for therapeutic modulation of Notch signaling in chronic disease settings.

More broadly, our findings support an emerging view that non-neuronal cells are integral components of hypothalamic metabolic circuitry. Understanding how glial state transitions influence neuronal computation and autonomic output may provide new therapeutic opportunities for obesity and metabolic disorders.

## ACKNOWLEDGEMENTS

We thank the Genentech Research Pathology, Necropsy, Histology and Next-Generation Sequencing labs for their experimental contributions. We thank Honglin Chen and Annalisa Lattanzi for their help with the design and order of AAV constructs. We thank Enric Llorens (Karolinska Intitutet) for feedback on the project. We appreciate the insightful feedback and comments on the paper from Heinrich Jasper and Fred de Sauvage. There were no external funding sources for this work.

## AUTHOR CONTRIBUTIONS

S.H. performed most experiments described below, analyzed and discussed data throughout the manuscript. M.H. performed some experiments, analyzed and discussed data throughout the manuscript. M.P.V. and G.F. analyzed and discussed the snRNA-seq. M.D. performed the single cell dissociation. D.S performed the metabolomics analysis. K.L., P.K., R.P., K.S, K.B., and A.S. provided experimental assistance. M.S and P.C. provided the PK analysis. B.J., A.ES., D.H., C.S., and C.H. provided intellectual feedback and guidance. L.M. designed and supervised the study, performed some experiments, provided feedback on all experimental design and data interpretation, and wrote the manuscript. All authors discussed the results and commented on the manuscript.

## MATERIALS AND METHODS

### Animal Models and Treatments

Animal experimentation at Genentech was performed according to protocols approved by the Genentech Institutional Animal Care and Use Program (IACUC) committee, in accordance with guidelines that adhere to and exceed state and national ethical regulations for animal care and use in research. Male C57BL/6 mice were obtained from Jackson Laboratories (strain ID: 00664). Ob/Ob mice were obtained from Jackson (strain ID: 00632) and PPARα-null (strain ID: 008154). *Notch1^floxed/floxed^* (Jackson 006951) and *Notch2^floxed/floxed^* (Jackson 010525) mice were crossed to generate *Notch1^floxed/floxed^ /Notch2^floxed/floxed^* mice. Fos-CreERT2 mice (Jackson 030323) were licensed from Stanford University and were crossed with LSL-tdTomato mice (Jackson 007904), licensed from Allen Institute for Brain Science. Male mice were injected with a combination of anti-Jag1 and anti-Jag2 antibodies or isotype controls. Four days after the antibodies’ injection, Fos-CreERT2;LSL-tdTomato mice were gavaged with 4-hydroxytamoxifen (Sigma, H6278, 40MPK) and sacrificed fifteen days post antibody injection. All mice were housed under specific pathogen-free (SPF) conditions and only males were used. Mice injected with blocking antibodies were 10-14 weeks of age, except for experiments in aged mice (>20 months). *Notch1^floxed/floxed^ /Notch2^floxed/floxed^*mice were 7-9 months old. Mice were injected intraperitoneally with a single dose of blocking antibodies diluted in PBS at the following concentrations: anti-Jag1 (YW167.71.70, 20MPK); anti-Jag2, (YW329.67.33, 20MPK)^13^; a combination of anti-Jag1 + anti-Jag2 (20MPK each); anti-Jag1/2 b50 (YW167.35.50, 15MPK), anti-Notch1 (YW169.60.79, 5MPK) and anti-Notch2 (YW167.54.76.76, 10MPK)^14^. Anti-Ragweed or anti-gD isotype control antibodies were injected at concentrations to achieve equal dose of the combination of antibodies in each study.

### Stereotaxic injection

*Notch1^floxed/floxed^/Notch2^floxed/floxed^* mice were bilaterally injected with adeno-associated virus to the hypothalamus in a similar manner to what has been previously described^39^. Animals were anesthetized with isoflurane in an induction chamber, then transferred to a stereotaxic frame (Stoelting Co.) where they were given continuous isoflurane via nose cone. A heating pad was used during surgery to maintain body temperature. Ophthalmic ointment was applied to both eyes before starting surgery. Meloxicam was administered for pain relief. The head was aligned on the stereotaxic frame using the earbars. Following this, a midline incision was made and the scalp was opened to identify bregma. A microdill with a burr tip 0.5 mm in diameter was used to make a small hole through the skull above the injection site on the left side of the head at -0.35 mm lateral from the midline and 1.32 mm caudal from bregma. A 10 μL Hamilton syringe (Hamilton, 701RN) with a 33 gauge needle (Hamilton, 7762-03) was slowly inserted to a depth of 5.65 mm. The needle was allowed to rest for 2 minutes, before infusing 1.5 μL of virus at a rate of 0.5 μL/minute. The needle was kept in place for at least 5 minutes post-injection to prevent backflow then slowly withdrawn. The same process is repeated on the right side of the skull. Once both bilateral injections have been made, the scalp incision is sealed with tissue adhesive (vetbond) and animals are allowed to recover on a heating pad.

### Viral Vectors

Recombinant AAV.Oligo1 viruses were produced in HEK293T cells by GenScript following standard triple plasmid transfection at 1:1:1 molar ratio (pHelper: pRepCap: pX552 transfer plasmid). The RepCap packaging plasmid contained the Oligo capsid sequence^44^. The AAV.Oligo1-Pdgfrα-Cre-T2A-BFP virus contains a payload driven by the tissue-specific Pdgfrα promoter^45^ to achieve tissue-specific expression in oligodendrocytes. The control virus AAV.Oligo-CAG-BFP-T2A-miRFP670nano3 contains a payload driven by a ubiquitous cytomegalovirus (CMV) enhancer/chicken β-actin hybrid (CAG) promoter. Expression of BFP upon AAV transduction allowed us to evaluate AAV transduction efficiency using RNAscope. Viruses were collected 72 hours post-transfection and purified by iodixanol gradient ultracentrifugation following the GenScript standard purification workflow. AAV plasmid constructs were synthesized *de novo* by GenScript.

### Micro-Computed Tomography (micro-CT)

The micro-CT imaging and image analysis technique to quantify body composition has been previously described in detail^46^. Briefly, animals were scanned under isoflurane anesthesia on an *in-vivo* µCT system (eXplore CT120, Trifoil Imaging, Chatsworth, CA, USA), Locus II, version 2.0.8.16 (GE Healthcare). Scan parameters were: 70 kV, 40 mA, 20 ms exposure, 900 views, 4×4 detector bin mode, 100 μm voxel size. An automated image analysis algorithm, based on intensity thresholding and morphological filtering, was used to segment the whole-body images into adipose tissue, lean tissue and bone^46^.

### Metabolic assays with animals

For the experiments assessing a high fat diet, we used the 60% Kcal from Fat formula (Envigo, TD. 06414). To measure food and water intake, as well as the amount of urine and feces excreted, we used the metabolic cages MM100 (Hatteras Instruments). Mice were singly housed in the metabolic cages 1 week before the experiment for acclimation purposes. The measurements and sample collection were taken every day at the same time. Blood glucose levels were measured in tip tail blood from living mice using a Contour glucometer and strips. For Glucose Tolerance Test (GTT), mice were fasted for 6 hours and 2MPK of 50%, Dextrose (NDC:0409-6648-02) diluted in PBS was injected intraperitoneally. Measurements of oxygen consumption, carbon dioxide production and mobility were taken every 15 min using Oxymax indirect calorimeter chambers (Columbus Instruments, 120455 and 110113). Measurements were collected using Clams Comprehensive Lab Animal monitoring system and analyzed using CLAMS data eXamination (CLAX) software. Animals were acclimatized for 6 days before and followed for up to 9 days after the treatment. For the cold challenge experiment, we used the Columbus cabinets (IT79SD) and mice were acclimatized to 4°C or 30°C for a week prior to the antibody treatments. Food intake in Figure 2 and for paired-feeding experiments was measured manually with a scale, while food intake in Figure 6 was measured automatically by the Oxymax Columbus system.

### Blood and serum analysis

Blood was collected at the time of necropsy. Blood cell counts were performed using the Sysmex XT-2000iV, an automated hematology analyzer for animal blood, which analyses white blood cells and reticulocytes with an optical detector block based on the flow cytometry method, using a semiconductor laser. It analyzes and outputs the results of 30 parameters of a blood sample. Red blood cells (RBCs) and platelet counts are analyzed by the RBC detector using the Hydro Dynamic Focusing method. Hemoglobin (HGB) is analyzed by the HGB detector based on the sodium lauryl sulphate hemoglobin detection method. For serum analysis, blood was collected in Microtainer® tubes (#365967 BD®) and centrifuged for 10 min at 10g to isolate the serum. Serum biochemistry was analyzed by Beckman AU480 using Beckman Coulter kits: Glucose (OSR6221), Cholesterol (OSR6216) and Triglycerides (OSR61118). The analysis of hormone levels in serums was performed with ELISA kits following the manufacturer protocol: Insulin (Crystal Chem, 90080), Glucagon (Crystal Chem, 81518) and Leptin (Abcam, ab199082). Cytokines in serum were analyzed using Milliplex MAP reagents (MilliporeSigma, St. Louis, MO) according to the manufacturer’s recommended protocol. Dilution of reconstituted standards were made at a 2.73-fold, to increase the number of data points from six to nine while maintaining original dynamic range. Fluorescence intensity was measured by xPONENT software v 4.2 on FlexMap 3D instruments (Luminex Corp, Austin, TX). Bio-Plex Manager v 6.2 (Bio-Rad Laboratories, Hercules, CA) was used to construct standard curves for each analyte on each plate using the median FI of at least 20 beads done in duplicate. A 4 or 5-point regression analysis was used to calculate best fit and cytokine levels.

### Histopathological stains and analysis

Tissue weights (brown adipose tissue, liver, etc.) were recorded postmortem. Tissue samples were collected at the time of necropsy and fixed in in 10% neutral buffered formalin (4% formaldehyde in solution), paraffin-embedded and cut in 4 µm sections, which were mounted in superfrost®plus slides and dried. Serial sections were stained with hematoxylin and eosin (HE) or Alcian Blue (Millipore, TMS010C) followed by nuclear fast red counterstaining (Vector, H3403). Immunohistochemistry was performed manually using standard techniques. In brief, sections were deparaffinized in xylene, rehydrated in ethanol and antigen retrieval was performed in DAKO target retrieval solution (#2369) for 15min. Endogenous peroxidase was blocked with 3% H2O2 for 20 min and slides were blocked in 5%BSA in PBS 0.1% Triton for 1 hour. Overnight incubation at room temperature (RT) was performed with primary antibodies diluted in blocking buffer as follows: anti-human IgG (1:500, Abcam, ab109489). Slides were incubated with Secondary antibody powervision poly-HRP anti-rabbit IgG (Leica PV6119) for 1 hour, and developed with TSA plus Cyanine 3 kit (Akoya Biosciences, NEL744001KT). Counterstaining with hematoxylin (1:10) was performed and slides were dehydrated with gradual ethanol and xylenes. Finally, slides were mounted with Permount for microscopic evaluation. RNAscope analysis was performed manually using the chromogenic RNAscopeTM 2.5 HD assay red kit (ACD, 322350) or RNAscope® 2.5 HD Duplex Reagent Kit (ACD, 322430) with the following transcript probes *Nrarp* (ACD, 411771), *Hes1* (ACD, 417701), *TagBFP* (ACD, 537141-C2), *Olig1* (ACD, 480651) and *Hes5* (ACD, 400991-C2) on formalin fixed paraffin-embedded liver, small intestines or brain sections. The antigen retrieval and protease-plus treatment were adjusted to 15 minutes. Quantification of brown adipose tissue (BAT) activity was done blindly, by assessing brown adipocyte size and accumulation of lipid vacuoles in 5 1 mm x 1 mm H&E sections per mouse, and averaging the scores. Score of 1 is white adipose tissue phenotype and score of 10 is BAT at its maximum capacity. Quantification of white adipose tissue (WAT) was performed using QuPath to analyze H&E sections^47^. The analysis followed a previously described system for quantifying adipocyte cell size and number^48^. A pixel classifier was trained to distinguish between cell membranes and lipid vacuoles. This was used on 1 mm x 1mm sections of epididymis WAT and inguinal region WAT, 3 ROIs per tissue. After the pixels were classified, new annotations were made with a minimum size of 115 μm and a minimum hole size of 30 μm. Annotations were manually confirmed to be only single adipocytes, then all annotations were measured. Size and number of adipocytes was plotted per animal.

### Whole-Brain Tissue Clearing

To visualize neuronal activation patterns with high anatomical fidelity, we employed the SHIELD (Stabilization to Harsh Conditions via Intramolecular Epoxide Linkage to prevent Degradation) protocol^49^. This approach was selected to stabilize endogenous cFos-tdTomato fluorescence against the harsh reagents required for clearing. Fixed brains were first incubated in SHIELD-OFF solution at 4°C for 3 days with mild shaking to ensure uniform penetration of the epoxide monomers. Samples were then transferred to SHIELD-ON solution and incubated at 37°C for 24 hours to complete the intramolecular cross-linking. Subsequent delipidation was performed in the SHIELD Delipidation Buffer for 7 days at 37°C. Finally, samples were immersed in EasyIndex at 37°C for 2 days until the refractive index (RI) was matched and the tissue reached complete transparency. A total of eight adult mice expressing the cFos-tdTomato reporter were used in this study. The subjects were divided into two experimental cohorts: Control (n = 4) and anti-Jag1/2-treated (n = 4).

### Whole-Brain 3D Lightsheet fluorescence microscopy and cFOS-tdTomato quantification

Volumetric imaging of SHIELD cleared brains was performed using a LaVision BioTec UltraMicroscope II (Blaze) light-sheet system. We utilized a 4x objective in combination with a 1.6x zoom lens (effective magnification=6.4). The tdTomato signal was acquired using a 561 nm laser. To ensure quantitative reproducibility across the anti-RW and anti-Jag1/2 groups, all acquisition parameters including laser power, sheet width, and exposure times were kept identical across all experimental samples. Raw unstitched 3D lightsheet TIFFs files were imported into BigStitcher (Fiji/ImageJ), where they were globally aligned via interest-point detection and fused into a single 16-bit 3D volume for downstream analysis. Following this, the fused TIFF files were further processed to remove horizontal shadow artifacts using PyStripe, a wavelet-based destriping algorithm. Standardized 3D analysis was conducted using the BrainGlobe computational suite^50^. The fused 561 nm channel was registered to the Allen Mouse Brain Common Coordinate Framework (CCFv3) at a 10 micron using Brainreg^51^. This registration provided the spatial transformation necessary to map individual samples into a common anatomical space. Within the same ecosystem, Cellfinder was utilized for the automated detection and segmentation of cFos-tdTomato+ nuclei^52^. To optimize detection for our specific imaging conditions, the cellfinder deep-learning model was retrained on a manually annotated subset of our data. Following the extraction of cell coordinates via cellfinder, data were processed using custom-written Python scripts to generate spatial heatmaps of neuronal activation. These scripts mapped the validated cFos+ centroids onto the CCFv3 template, applying a Gaussian kernel to visualize activation “hotspots” and density gradients across the 3D brain volume^53^. These heatmaps were used to qualitatively assess the differences in activation patterns between anti-RW and anti-Jag1/2-treated brains. Quantitative outputs were generated as cell counts and densities cells/mm^3^) for each of the hierarchical regions defined by the Allen Atlas.

### Global Metabolomics Analysis

Hypothalamus tissues (20-40 mg) were homogenized in 375 μL of ice cold 80:20 Methanol: Water containing in-house metabolomics and lipidomics Recovery IS (Mixture of Stable Isotope Labeled Internal Standards) using a bead beater homogenizer followed by centrifuging at 4000 RPM for 5 min at 4°C after a 30 min incubation period in the -20°C freezer. Supernatant was aliquoted into a separate tube. To the sample homogenate pellet, 300 μL of cold chloroform were added again followed by vortex mixing thoroughly and centrifuging at 4000 RPM for 10 min at 4°C. Supernatant was aliquoted and combined into the above tube with previous supernatant followed by addition of 195 μL of ice-cold water, vortex mixing and centrifuging at above conditions for phase separation and liquid-liquid extraction. 300 µL of top layer (MeOH/water) were used for Global small molecule polar and semi-polar Metabolomics analysis. Metabolomics sample aliquots were dried under nitrogen and reconstituted with 90 µL of 8:2 Acetonitrile:Water and 10 µL of Global metabolomics stable isotope labeled internal standards mix followed by shaking at 800 RPM for 5 min and 5 μL were injected for HILIC MS analysis as described below. Pool QCs were also prepared as run quality control samples by combining a small volume (20 μL) after phase separation. Cell pellets were homogenized by sonication for 10 min in 500 μL of ice cold 80:20 Methanol: Water containing in-house metabolomics and lipidomics Recovery IS and centrifuged at 4°C for 1 minute. 375 μL of supernatant were aliquoted into a separate tube and the rest of the process is similar to tissue sample preparation. 100 μL of media samples were precipitated with 375 μL of ice cold 80:20 Methanol:Water containing in-house metabolomics and lipidomics Recovery IS and rest of the sample preparation was similar to tissue samples. For global metabolomics, ACQUITY UPLC BEH Amide Column (2.1 mm × 150 mm × 1.7 μm, 130 Å, Waters Corporation) was used to separate metabolites with mobile phase A of 100% water containing 10 mM ammonium formate and 0.125% formic acid, and mobile phase B of 95% acetonitrile in water containing 10 mM ammonium formate and 0.125% formic acid. Data acquisition was achieved on a Shimadzu Nexera HPLC series system (Shimadzu, Japan) coupled with a Thermo Q Exactive Plus Hybrid Quadrupole-Orbitrap Mass Spectrometer (Thermo Fisher Scientific). Injection volume of 5 μL was used for sample analysis under Heated Electrospray Ionization (HESI) condition with detailed instrument parameters described previously^54^. Metabolites were identified at Level 1 confidence by matching at least two independent orthogonal experimental data (accurate mass, retention time) against compound retention time library published previously.^55^ Trend analysis of stable isotope labeled internal standards and matrix PoolQC samples were examined (with %RSD less than 50%) to validate system suitability and data robustness. For each metabolite, relative quantification was obtained through either MS peak area of analyte or MS peak area ratio of analyte/internal standard using inhouse data processing tools based on XCMS^56^.

### Targeted Metabolomics Analysis

Tissue, cells homogenates and media samples were also analyzed for targeted metabolomics analysis of TCA cycle metabolites (Citrate, Isocitrate, Lactate, Succinate, Pyruvate, Fumarate, Malate, alpha-ketoglutarate, alpha-hydroxyglutarate, hydroxy-butyrate) and amino acids (Asparate, Glutamate, Glutamine, N-methyl glutamate and N-methyl Asparate). TCA cycle intermediates targeted quantitative analysis was conducted as described previously^57^. Briefly, analytes were derivatized using 1M O-BHA and 1M EDC in pyridine-HCl buffer followed by liquid liquid extraction with ethyl acetate and Liquid chromatography high resolution mass spectrometry (LC-HRMS) full scan (100-900 m/z, positive mode) analysis on a ThermoFisher Q-Exactive Exploris high resolution mass spectrometer with a HESI-II ion source (Thermo Scientific, Waltham, MA, USA). Source parameters were adjusted to flow rate and liquid chromatographic parameters as described previously^57^. Targeted TCA panel quantitative analysis included calibration curves using metabolite standards in water and stable isotope labeled internal standards for each metabolite to account for analyte recovery, derivatization efficiency and ion suppression as described previously^57,58^. For targeted Amino acids analysis above samples were processed using derivatization using PITC as described previously^59^. Briefly, samples and calibrators were derivatized using 50 μL of 5% PITC derivatization solution (300 μL of PITC reagent + 1900μL each of ethanol, water and pyridine). LC-HRMS full scan analysis was conducted for targeted quantitation of Aminoacid metabolites. Stable isotope labeled internal standards for Glutamate, Aspartate and Glutamine were included in the analysis to account for analyte recovery, derivatization efficiency, and ion suppression.

### Serum metabolomics

For serum analysis, blood was collected in Microtainer® tubes (BD Biosciences, 365967) and centrifuged for 10 min at 10 g to isolate the serum. For the Tricarboxylic acid (TCA) cycle panel, a derivatization assay was performed with method details previously reported^57^. Derivatization of the tricarboxylic acid intermediates with O-benzylhydroxylamine for liquid chromatography-tandem mass spectrometry detection^60^. Anal Biochem. 2014]. Data acquisition was achieved on a Shimadzu Nexera HPLC series system (Shimadzu, Kyoto, Japan) coupled with a Sciex 6500 Mass Spectrometer (Sciex, Framingham, MA, USA). Injection volume of 7.5 µL was used for sample analysis under Electrospray Ionization (ESI) condition. Samples were run under positive mode. The Sciex 6500 Mass Spectrometer was operated with the following parameters: Temperature, 500 °C; Gas 1, 60 units; Gas 2, 60 units; IS, 5500V; Declustering potential, 100V; EP, 10V; CXP, 10V. For targeted metabolomics, absolute quantification (concentration) was obtained using MultiQuant software (SCIEX, Framingham, MA, USA) using calibration curves for each TCA cycle metabolite. For glycolysis analysis, all samples, standards, blanks, and QCs were diluted 1:1 with water. A 20 µL aliquot was mixed with 250 µL cold methanol and vortexed. 750 µL Methyl tert-butyl ether was then added, and all the samples were vortexed once more. Afterwards, 200 µL water was added to all samples and vortexed before the bottom layer was taken out and dried under nitrogen gas at 37°C for one hour. These samples were then reconstituted with 100 µL water before being injected by LC-MS/MS. Data acquisition and quantitation were performed using the same instrumentation and software as the TCA analysis. Injection volume of 7.5 µL was used for sample analysis under Electrospray Ionization condition. Samples were run under negative mode. The Sciex 6500 Mass Spectrometer was operated with the following parameters: Curtain gas, 30; Temperature, 500 °C; Gas 1, 60 units; Gas 2, 50 units; IS, -4500V; EP, -10V.

### PK Analysis

Lungs, brains (which were dissected to isolate hypothalamus), and serum were collected 1, 2, 3, and 7 days after anti-Jag1 and anti-Jag2 antibody treatment. Lung and brain samples were lysed using 1% NP40 in Tissue Lyser II (Qiagen). Nunc® MaxiSorp™ 384-well plates (Thermo, cat# 464718) were coated with 1 μg/mL monkey adsorbed sheep anti-Human IgG (Binding Site, cat#AU003.M) diluted in 0.05 M carbonate/bicarbonate buffer pH 9.6 and incubated overnight at 4°C. The plates were washed 3 times with a wash buffer (0.05% Tween-20 in PBS buffer, pH 7.4) and treated with block buffer (PBS/0.5% BSA/15 ppm Proclin, pH 7.4) for 1 to 2 hours at room temperature (RT). The plates were washed 3 times with wash buffer and then samples diluted in sample diluent (PBS/0.5% BSA/0.05% Tween 20/5mM EDTA/0.25% CHAPS/ 0.35M NaCl/15 ppm Proclin, pH 7.4) were added to the wells and incubated for 2 hours at RT with gentle agitation. After washing the plates 6 times with wash buffer, a detection antibody, HRP-conjugated F(ab′)2 goat anti-human IgG and Fc-specific polyclonal antibody (Jackson ImmunoResearch, 109-036-098), diluted to 12.5 ng/mL in assay buffer (PBS/0.5% BSA/15 ppm Proclin/0.05% Tween 20, pH7.4) was added to the wells and incubated on a shaker for 1 hour at RT. The plates were washed 6 times with a wash buffer and developed using TMB peroxidase substrate (KPL, 5120-0077) for 20 minutes followed by 1 M Phosphoric acid to stop the reaction. Absorbance was measured at 450 nm against a reference wavelength of 620 nm. The concentration of the samples was extrapolated from a 4-parameter fit of the standard curve.

Please note: for coat, detection, TMB, Acid - 25 µL volume added to each 384 well; for block 50 µL volume added to each 384 well. Different lots of antibodies may affect optimal final concentration in the assay.

### Flow Cytometry

For FACS analysis of oligodendrocytes, brain tissues were dissociated using the Adult Brain Dissociation kit (Miltenyi; 130-107-677) according to the manufacturer’s protocol. The resulting single-cell suspension was then stained extracellularly with an antibody cocktail for 20 minutes on ice. Subsequently, samples were fixed and permeabilized using the eBioscience™ Foxp3/Transcription Factor Staining Buffer Set (Thermo Fisher Scientific; 00-5523-00) for 30 min at RT. Intracellular staining with an antibody cocktail was performed at 4°C overnight. Stains were performed with the following antibodies: anti-CD45 (Invitrogen; 58-0451-82), anti-Myelin Basic Protein (MBP, Biolegend; 850905), anti-PDGFRα (Biolegend; 135911), anti-Olig2 (Cell Signaling Technology; 43829), and Live/dead NIR (ThermoFisher Scientific; L34975). Flow cytometry data was collected via Cytek^®^ Aurora and analyzed using FlowJoV10.

### Luciferase reporter Assay

U87 glioblastoma cells which endogenously express predominantly Notch2, but only very low levels of other Notch receptors were co-transfected with a Notch-responsive TP-1 (12X CSL) Firefly luciferase reporter and a constitutively expressed Renilla luciferase reporter (pRL-CMV, Promega #E2261) to control for transfection efficiency. Antibodies, DAPT (5 µM) or DMSO vehicle control were added with the ligand-expressing cells (NIH-3T3 cells stably transfected with human Jag1 or Jag2) six to eight hours after transfection. Luciferase activities were measured after 20 hours of co-culture (Dual Glo Luciferase, Promega, E2920), using a Perkin-Elmer EnVison 2103 Multilabel Reader. Typically, four replicates are analyzed for each condition, and values are expressed as relative luciferase units (Firefly signal divided by the Renilla signal) and graphed as % activation per DMSO control.

### *In Vitro* Differentiation Assays

Primary astrocytes were cultured from mouse pups on postnatal day 1-3 as previously described^61,62^. Briefly, cortices were isolated with meninges removed, tissue was trypsinized in 0.5% trypsin (Gibco; 15400054) for 10 minutes, then filtered through a 70 micron filter, spun at 700 g for 5 min, then resuspended in 40 mL of DMEM containing 10% FBS, 1% glutamax (Gibco; 35050061), and 1% penicillin streptomycin (Gibco; 15140122). Cell suspension was cultured in a T225 flask for 3 days, washed with PBS three times, then the media was replaced and cells were allowed to grow until 10-14 days after isolation. Microglia were shaken off and Astrocytes were trypsinized and replated for co-culture assays. Primary OPCs were cultured from mouse pups on postnatal days 5-7 as previously described^63^. Briefly, cortices were isolated with meninges removed and digested in papain (Worthington; LS003118) for 45-60 min at 34℃ with 5% CO_2_. Cells were filtered with Ovo (Worthington; LS003086) gradients then immuno-panned to select for purity. First non-specific debris was removed with a 60 min incubation on an anti-BSL-1 (VectorLabs; L1100) coated plate and then OPCs were enriched by incubating cells on plates coated with anti-CD104a (BD Pharmigen; 558774). Cells were released from the plate with gentle trypsinization and replated onto a bed of astrocytes.

Co-cultures were grown in ratios of roughly 1:3 astrocytes:OPCs with about 15,000 astrocyte cells plated/well and 45,000 OPC cells plated/well in 96-well plates. Co-cultures grew together for 2-7 days then were treated for a given number of days with anti-RW or anti-Notch1 (YW169.60.79) and anti-Notch2 (YW167.54.76.76) antibodies 10 µg/mL. Cells were fixed in 4% paraformaldehyde and stained for myelin basic protein (Abcam; ab62631, 1:500), olig2 (ThermoFisher, OSO00005W; 1:300), and GFAP (Abcam; ab4674, 1:600). Images were taken using a Leica THUNDER imaging system and analyzed using ImageJ. 9 images were taken per well and data is represented as fold change compared to the average of anti-RW-treated cells.

### RNA isolation and bulk RNAseq

Total RNA was extracted from snap-frozen liver and hypothalamus tissues with Trizol (Invitrogen) following the manufacturer’s recommendations. After adding isopropanol, the samples were transferred into the RNAeasy extraction column (Qiagen) and we washed and eluted following the manufacturer protocol. For bulk RNAseq, RNA concentrations and quality were assessed using the Qubit RNA HS Assay Kit (Thermo Fisher Scientific) and RNA ScreenTape on TapeStation 4200 (Agilent Technologies). cDNA libraries were generated from 2 ng of total RNA using the Smart-Seq V4 Ultra Low Input RNA Kit (Takara). 150 pg of cDNA was used to make sequencing libraries with the Nextera XT DNA Sample Preparation Kit (Illumina). Libraries were quantified with the Qubit dsDNA HS Assay Kit (Thermo Fisher Scientific), and the average library size was determined using D1000 ScreenTape on TapeStation 4200 (Agilent Technologies). Libraries were pooled and sequenced on HiSeq 4000 (Illumina) to generate 30-40 million single-end 50-base pair reads for each sample. Sequencing reads were aligned to the mouse genome (NCBI Build 38) using GSNAP^75^ version ‘2013-10-10’, allowing a maximum of two mismatches per 50 base pair sequence (parameters: ‘-M 2 -n 10 -B 2 -i 1 -N 1 -w 200000 -E 1 --pairmax-rna=200000 --clip-overlap’). Expression levels were quantified by calculating the number of reads mapped to the exons of each RefSeq gene using the HTSeqGenie R package. Transcript annotation relied on the RefSeq database NCBI Annotation Releases 104 for mouse. Read counts were processed with DeSeq2 (v1.48.2)^64^ and scaled using variance stabilizing transformation. Differential expression analysis was performed on the counts using the Wald test with Benjamini & Hochberg correction. Gene set enrichment analysis was performed with fgsea (v1.34.2)^65,66^ using the CP:REACTOME gene set from the molecular signature database. Where indicated, gene expression was normalized as Reads Per Kilobase gene model per Million total reads (nRPKM).

### Hypothalamus Single-Cell Nuclei isolation and library prep

Nuclei for single-nucleus sequencing from ad libitum fed mice were isolated as described previously^67^. Briefly, mice were euthanized by CO_2_ inhalation and decapitation. The brain was quickly excised intact and placed on ice. Under a dissection scope, the hypothalamus was dissected and placed in 1mL ice-cold lysis buffer (20mM NaCl,5mM MgCl2, 0.1% TX-100, 10mM Tris–HCl pH 7.2) containing EDTA Free protease inhibitor (Sigma, 4693124001), RNAse inhibitor 0.2 U/mL (Life Technologies, N8080119), and RNase-free DNase (Promega Corporation, M6101), in an 2 mL Dounce homogenizer tube (Kimble Chase, 885300-0002). Dounce homogenization was performed using 5-10 strokes with a “loose’ pestle to preserve the integrity of neuronal nuclei. The lysate was filtered through a 70-micron filter and centrifuged at 300 g for 5 min. The pellet was resuspended in 1 mL nuclei suspension buffer (NSB) (HBSS (Thermo Fisher, 14025092) containing 3% BSA Fraction VI and RNAse inhibitor). Nuclei were further purified using an Optiprep (Sigma, D-1556) density gradient (16%, 35% in HBSS). The above filtered lysate was mixed with an equal volume of 16% optiprep and placed carefully over the gradient in a 15 mL tube. The gradients were centrifuged in a swinging bucket rotor at 2500 g for 20 min. The purified nuclei were then carefully aspirated and resuspended in NSB for another 300 g centrifugation to remove the residual Optiprep. Nuclei were labeled with propidium iodide and DAPI for isolation and counting by FACS using a FACSAria Fusion Flow Cytometer (BD Biosciences) before loading onto a 10X Genomics fluidics chip for droplet generation and barcoding. Nuclei from the hypothalamus of paired-fed mice were isolated using Chromium Nuclei Isolation Kit with RNase Inhibitor (10x Genomics, PN-1000494) following the manufacturer’s instructions. Before the last wash and centrifugation step anti-Nucleus MicroBeads (Miltenyi Biotec, 130-132-997) were added to the tissue lysate following the manufacturer’s instructions. Nuclei were resuspended in 200 µL Wash and Resuspension Buffer and loaded onto a 10X Genomics fluidics chip for droplet generation and barcoding (10X Genomics, PN-1000127).

Single nuclei expression libraries from ad libitum and paired-fed mice were prepared using the Chromium Next GEM Single Cell 3ʹ GEM, Library & Gel Bead Kit v3.1 (10X Genomics, PN-1000121) according to the manufacturer’s instructions. Libraries were quantified and profiled using a Bioanalyzer High Sensitivity DNA kit (Agilent Technologies, 5067-4626). Libraries were sequenced by HiSeq4000 (Illumina) following the 10X Genomics sequencing specifications.

### Hypothalamus Single-Cell Nuclei Analysis

Data was processed with cellranger-7.1.0 using the GRCm38 transcriptome and CellBender (v0.3.0) to reduce signal coming from ambient RNA. Data processing was done in R (v4.5.0). After filtering for high quality cells (nFeature_RNA > 200 & nFeature_RNA < 5000 & percent.mt <10), downstream analysis was performed in Seurat (v5.3.0)^68^, where 30 clusters were identified using default FindCluster parameters (resolution = 0.8, algorithm = 1). Dimension reduction was performed using UMAP on the first 20 dimensions. Labeltransfer was performed with FindTransferAnchors using the top 30 PCA dimensions of the HypoMap dataset at two granularities using the Author_Class and C66_named meta data columns^27^. Differential abundance analysis was performed with MiloR (v2.4.1)^28^ using parameters k = 20, d = 10. To prioritize transcriptomic response of cell lineages, Augur (v1.0.3)^38^ analysis was performed on the top 5000 highly variable genes using the random forest classifier (n_subsamples = 100, trees = 500). Differentially expression analysis was done using FindMarkers, and significant genes (p_val< 0.01, avg_log2FC > 0.5) were visualized with EnhancedVolcano or (v1.26.0) fed into gene ontology analysis using enrichR (3.4)^69^ on the GO_Biological_Process_2025 database. To investigate the oligodendrocyte lineage with higher granularity, UMAP dimension reduction was performed solely on these cells (dims = 1:10, min.dist = 0.4) and cells were re-clustered (resolution = 0.2). Trajectory inference was performed with slingshot (v2.16.0) using default parameters. Cell-communication analysis was inferred with CellChat (v2.2.0)^37^ based on the non-protein signaling database, enriched for genes related to neuron-neuron and metabolic communication. Fasting signatures were derived from the HypoMap by comparing different neuronal populations across normal chow vs. fasted conditions (p_val_adj < 0.01, avg_log2FC > 1), and evaluated in data from this study using AddModuleScore.

### Statistical analysis

All values in graphs are presented as mean ± s.e.m. Replicates are described in the figure legends. Mice were randomly assigned to treatment groups for *in vivo* studies. Significant differences between two groups were evaluated using a two-tailed unpaired Welch’s *t*-test or Mann-Whitney test. Multiple unpaired two-stage step-up (Benjamini, Krieger, and Yekutieli) *t*-test or non-parametric Mann-Whitney test were used for comparisons between more than three parameters. Two-way ANOVA or Mixed-effects analysis followed by multiple comparisons were used to analyze two groups with repeated measures over time. Statistical significance was noted as (\**p* < 0.05, \*\**p* < 0.01, \*\*\**p* < 0.001, \*\*\*\**p* < 0.0001).

## DATA AVAILABILITY

All RNA-seq data and single-nucleus RNA-seq data will be made publicly available as of the date of this publication.

## CODE AVAILABILITY

This paper does not report original code. Any additional information required to reanalyze the data reported in this paper is available at github.com/mpverhagen/Notch-hypothalamus.

“Extended Data is available for this paper.”

## REFERENCES

1. Fruh, S. M. Obesity: Risk factors, complications, and strategies for sustainable long-term weight management. J. Am. Assoc. Nurse Pract. 29, S3–S14 (2017).

2. Seoane-Collazo, P. et al. Hypothalamic-autonomic control of energy homeostasis. Endocrine 50, 276–291 (2015).

3. Aponte, Y., Atasoy, D. & Sternson, S. M. AGRP neurons are sufficient to orchestrate feeding behavior rapidly and without training. Nat. Neurosci. 14, 351–355 (2011).

4. Millington, G. W. The role of proopiomelanocortin (POMC) neurons in feeding behaviour. Nutr. Metab. 4, 18 (2007).

5. Deem, J. D., Faber, C. L. & Morton, G. J. AgRP neurons: Regulators of feeding, energy expenditure, and behavior. FEBS J. 289, 2362–2381 (2022).

6. Nampoothiri, S., Nogueiras, R., Schwaninger, M. & Prevot, V. Glial cells as integrators of peripheral and central signals in the regulation of energy homeostasis. Nat. Metab. 4, 813–825 (2022).

7. Djogo, T. et al. Adult NG2-Glia Are Required for Median Eminence-Mediated Leptin Sensing and Body Weight Control. Cell Metab. 23, 797–810 (2016).

8. Buller, S. et al. Median eminence myelin continuously turns over in adult mice. Mol. Metab. 69, 101690 (2023).

9. Bray, S. J. & Bigas, A. Modes of Notch signalling in development and disease. Nat. Rev. Mol. Cell Biol. 26, 522–537 (2025).

10. Siebel, C. & Lendahl, U. Notch Signaling in Development, Tissue Homeostasis, and Disease. Physiol. Rev. 97, 1235–1294 (2017).

11. Wu, Y. et al. Therapeutic antibody targeting of individual Notch receptors. Nature 464, 1052– 1057 (2010).

12. Milano, J. et al. Modulation of Notch Processing by γ-Secretase Inhibitors Causes Intestinal Goblet Cell Metaplasia and Induction of Genes Known to Specify Gut Secretory Lineage Differentiation. Toxicol. Sci. 82, 341–358 (2004).

13. Lafkas, D. et al. Therapeutic antibodies reveal Notch control of transdifferentiation in the adult lung. Nature 528, 127–131 (2015).

14. Wu, Y. et al. Therapeutic antibody targeting of individual Notch receptors. Nature 464, 1052– 1057 (2010).

15. Ridgway, J. et al. Inhibition of Dll4 signalling inhibits tumour growth by deregulating angiogenesis. Nature 444, 1083–1087 (2006).

16. Tran, E. et al. Immune targeting of fibroblast activation protein triggers recognition of multipotent bone marrow stromal cells and cachexia. J. Exp. Med. 210, 1125–1135 (2013).

17. van Es, J. H. et al. Notch/γ-secretase inhibition turns proliferative cells in intestinal crypts and adenomas into goblet cells. Nature 435, 959–963 (2005).

18. Fre, S. et al. Notch signals control the fate of immature progenitor cells in the intestine. Nature 435, 964–968 (2005).

19. Schröder, N. & Gossler, A. Expression of Notch pathway components in fetal and adult mouse small intestine. Gene Expr. Patterns 2, 247–250 (2002).

20. Coleman, D. L. Obese and diabetes: Two mutant genes causing diabetes-obesity syndromes in mice. Diabetologia 14, 141–148 (1978).

21. Lee, S. S.-T. et al. Targeted Disruption of the α Isoform of the Peroxisome Proliferator-Activated Receptor Gene in Mice Results in Abolishment of the Pleiotropic Effects of Peroxisome Proliferators. Mol. Cell. Biol. 15, 3012–3022 (1995).

22. Mosteiro, L. et al. Notch signaling in thyrocytes is essential for adult thyroid function and mammalian homeostasis. Nat. Metab. 5, 2094–2110 (2023).

23. Yang, Y. et al. Caloric Restriction Remodels Energy Metabolic Pathways of Gut Microbiota and Promotes Host Autophagy. 2020.08.16.251215 Preprint at 10.1101/2020.08.16.251215 (2020).

24. Dankel, S. N. et al. Changes in Plasma Pyruvate and TCA Cycle Metabolites upon Increased Hepatic Fatty Acid Oxidation and Ketogenesis in Male Wistar Rats. Int. J. Mol. Sci. 24, 15536 (2023).

25. Ahima, R. S. et al. Role of leptin in the neuroendocrine response to fasting. Nature 382, 250– 252 (1996).

26. Mahoney, L. B., Denny, C. A. & Seyfried, T. N. Caloric restriction in C57BL/6J mice mimics therapeutic fasting in humans. Lipids Health Dis. 5, 13 (2006).

27. Steuernagel, L. et al. HypoMap—a unified single-cell gene expression atlas of the murine hypothalamus. Nat. Metab. 4, 1402–1419 (2022).

28. Dann, E., Henderson, N. C., Teichmann, S. A., Morgan, M. D. & Marioni, J. C. Differential abundance testing on single-cell data using k-nearest neighbor graphs. Nat. Biotechnol. 40, 245–253 (2022).

29. Wang, S. et al. Notch receptor activation inhibits oligodendrocyte differentiation. Neuron 21, 63–75 (1998).

30. John, G. R. et al. Multiple sclerosis: Re-expression of a developmental pathway that restricts oligodendrocyte maturation. Nat. Med. 8, 1115–1121 (2002).

31. Zhang, Y. et al. Notch1 signaling plays a role in regulating precursor differentiation during CNS remyelination. Proc. Natl. Acad. Sci. 106, 19162–19167 (2009).

32. Wang, C. et al. IL-17 induced NOTCH1 activation in oligodendrocyte progenitor cells enhances proliferation and inflammatory gene expression. Nat. Commun. 8, 15508 (2017).

33. Viganò, F. et al. GPR17 expressing NG2-Glia: Oligodendrocyte progenitors serving as a reserve pool after injury. Glia 64, 287–299 (2016).

34. Nave, K.-A., Asadollahi, E. & Sasmita, A. Expanding the function of oligodendrocytes to brain energy metabolism. Curr. Opin. Neurobiol. 83, 102782 (2023).

35. Philips, T. & Rothstein, J. D. Oligodendroglia: metabolic supporters of neurons. J. Clin. Invest. 127, 3271–3280 (2017).

36. Fünfschilling, U. et al. Glycolytic oligodendrocytes maintain myelin and long-term axonal integrity. Nature 485, 517–521 (2012).

37. Jin, S., Plikus, M. V. & Nie, Q. CellChat for systematic analysis of cell–cell communication from single-cell transcriptomics. Nat. Protoc. 20, 180–219 (2025).

38. Skinnider, M. A. et al. Cell type prioritization in single-cell data. Nat. Biotechnol. 39, 30–34 (2021).

39. Lau, J. et al. CART neurons in the arcuate nucleus and lateral hypothalamic area exert differential controls on energy homeostasis. Mol. Metab. 7, 102–118 (2018).

40. Allen, W. E. et al. Thirst-associated preoptic neurons encode an aversive motivational drive. Science 357, 1149–1155 (2017).

41. Drucker, D. J. Mechanisms of Action and Therapeutic Application of Glucagon-like Peptide-1. Cell Metab. 27, 740–756 (2018).

42. Kim, J. G. et al. Leptin signaling in astrocytes regulates hypothalamic neuronal circuits and feeding. Nat. Neurosci. 17, 908–910 (2014).

43. García-Cáceres, C. et al. Astrocytic Insulin Signaling Couples Brain Glucose Uptake with Nutrient Availability. Cell 166, 867–880 (2016).

44. Powell, S. K. et al. Characterization of a novel adeno-associated viral vector with preferential oligodendrocyte tropism. Gene Ther. 23, 807–814 (2016).

45. Nathanson, J. L. et al. Short Promoters in Viral Vectors Drive Selective Expression in Mammalian Inhibitory Neurons, but do not Restrict Activity to Specific Inhibitory Cell-Types. Front. Neural Circuits 3, 19 (2009).

46. Wyatt, S. K. et al. Fully-automated, high-throughput micro-computed tomography analysis of body composition enables therapeutic efficacy monitoring in preclinical models. Int. J. Obes. 39, 1630–1637 (2015).

47. Bankhead, P. et al. QuPath: Open source software for digital pathology image analysis. Sci. Rep. 7, 16878 (2017).

48. Palomäki, V. A., Koivukangas, V., Meriläinen, S., Lehenkari, P. & Karttunen, T. J. A Straightforward Method for Adipocyte Size and Count Analysis Using Open-source Software QuPath. Adipocyte 11, 99–107.

49. Park, Y.-G. et al. Protection of tissue physicochemical properties using polyfunctional crosslinkers. Nat. Biotechnol. 37, 73–83 (2019).

50. Claudi, F. et al. BrainGlobe Atlas API: a common interface for neuroanatomical atlases. J. Open Source Softw. 5, 2668 (2020).

51. Tyson, A. L. et al. Accurate determination of marker location within whole-brain microscopy images. Sci. Rep. 12, 867 (2022).

52. Tyson, A. L. et al. A deep learning algorithm for 3D cell detection in whole mouse brain image datasets. PLOS Comput. Biol. 17, e1009074 (2021).

53. Claudi, F. et al. Visualizing anatomically registered data with brainrender. eLife 10, e65751 (2021).

54. Browder, K. C. et al. In vivo partial reprogramming alters age-associated molecular changes during physiological aging in mice. *Nat*. Aging 2, 243–253 (2022).

55. Bonini, P., Kind, T., Tsugawa, H., Barupal, D. K. & Fiehn, O. Retip: Retention Time Prediction for Compound Annotation in Untargeted Metabolomics. Anal. Chem. 92, 7515–7522 (2020).

56. Louail, P. et al. xcms in Peak Form: Now Anchoring a Complete Metabolomics Data Preprocessing and Analysis Software Ecosystem. Anal. Chem. 97, 27639–27645.

57. Tan, B., Lu, Z., Dong, S., Zhao, G. & Kuo, M.-S. Derivatization of the tricarboxylic acid intermediates with *O*-benzylhydroxylamine for liquid chromatography–tandem mass spectrometry detection. Anal. Biochem. 465, 134–147 (2014).

58. Lai, Z. et al. LC-HRMS-based targeted metabolomics for high-throughput and quantitative analysis of 21 growth inhibition-related metabolites in Chinese hamster ovary cell fed-batch cultures. Biomed. Chromatogr. 36, e5348 (2022).

59. Zheng, J., Zhang, L., Johnson, M., Mandal, R. & Wishart, D. S. Comprehensive Targeted Metabolomic Assay for Urine Analysis. Anal. Chem. 92, 10627–10634 (2020).

60. Tan, B., Lu, Z., Dong, S., Zhao, G. & Kuo, M.-S. Derivatization of the tricarboxylic acid intermediates with *O*-benzylhydroxylamine for liquid chromatography–tandem mass spectrometry detection. Anal. Biochem. 465, 134–147 (2014).

61. Byrnes, A. E. et al. A fluorescent splice-switching mouse model enables high-throughput, sensitive quantification of antisense oligonucleotide delivery and activity. *Cell Rep*. Methods 4, 100673 (2024).

62. Schildge, S., Bohrer, C., Beck, K. & Schachtrup, C. Isolation and culture of mouse cortical astrocytes. J. Vis. Exp. JoVE 50079 (2013) doi:10.3791/50079.

63. Dugas, J. C. & Emery, B. Purification of oligodendrocyte precursor cells from rat cortices by immunopanning. Cold Spring Harb. Protoc. 2013, 745–758 (2013).

64. Love, M. I., Huber, W. & Anders, S. Moderated estimation of fold change and dispersion for RNA-seq data with DESeq2. Genome Biol. 15, 550 (2014).

65. Korotkevich, G. et al. Fast gene set enrichment analysis. 060012 Preprint at 10.1101/060012 (2021).

66. Subramanian, A. et al. Gene set enrichment analysis: A knowledge-based approach for interpreting genome-wide expression profiles. Proc. Natl. Acad. Sci. 102, 15545–15550 (2005).

67. Jung, M. et al. Cross-species transcriptomic atlas of dorsal root ganglia reveals species-specific programs for sensory function. Nat. Commun. 14, 366 (2023).

68. Hao, Y. et al. Dictionary learning for integrative, multimodal and scalable single-cell analysis. Nat. Biotechnol. 42, 293–304 (2024).

69. Chen, E. Y. et al. Enrichr: interactive and collaborative HTML5 gene list enrichment analysis tool. BMC Bioinformatics 14, 128 (2013).

