## Extended data for "Hypothalamic oligodendrocytes regulate systemic energy balance through Notch-dependent state transitions"

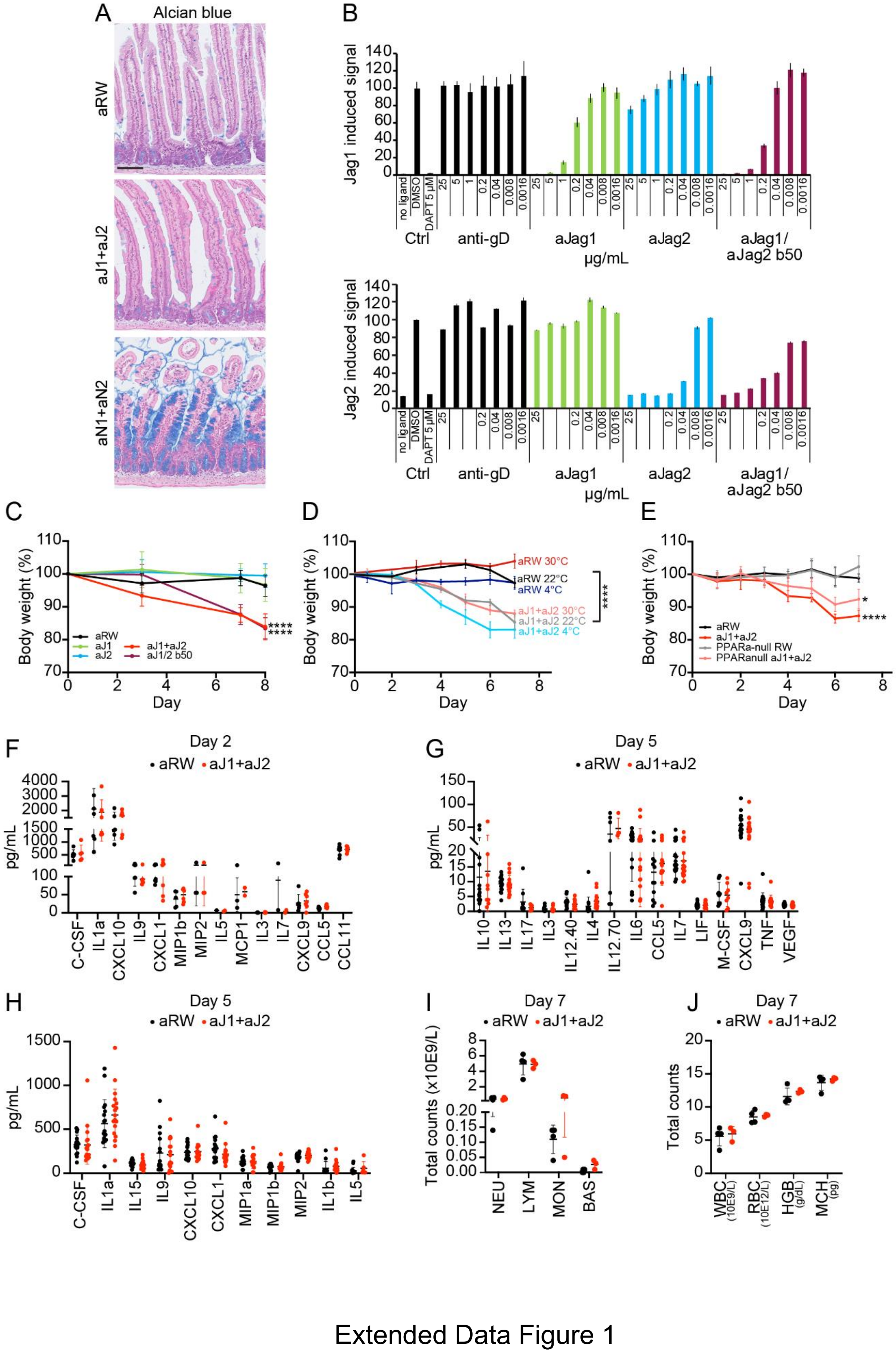


Extended Data Figure 1

**Extended Data Figure 1. Body weight reduction after anti-Jag1/2 treatment is independent of intestinal or hematological phenotypes or inflammation.**

A. Alcian blue staining in the intestines of mice treated with anti-Ragweed (anti-RW) isotype control antibody (RW); a combination of anti-Jag1 and anti-Jag2 blocking antibodies (anti-Jag1/2) or a combination of anti-Notch1 and anti-Notch2 blocking antibodies (anti-Notch1/2). The scale bar is 100 µm.

B. Luciferase reporter assay in U87 cell line. Y axis indicates luciferase activity represented as percent of inhibition compared to DMSO-treated condition (at 100%) and X axis different concentrations of the indicated antibodies (in µg/mL). All values are expressed as mean ± s.d. of n=4 replicates per group. Ctrl, controls.

C. Percentage of body weight compared to initial body weight over 8 days. Treatments were anti-RW antibody; anti-Jag1; anti-Jag2, anti-Jag1/2 or a dual anti-Jag1/2 b50. All values are expressed as mean ± s.d. of n=8 mice per group. Statistical significance was assessed using two-way ANOVA with Dunnett's multiple comparison: at day 8, $p<0.0001$, ****.

D. Percentage of body weight reduction compared to initial body weight in mice after 7 days of treatment at different ambient temperatures (22°C, 4°C or 30°C). Treatments were anti-RW or anti-Jag1/2. All values are expressed as mean ± s.d. and at least n=4 mice per group. Statistical significance was assessed using Mixed-effects analysis: $p<0.0001$, **** (source of variance is treatment).

E. Percentage of body weight compared to initial body weight in PPARa-null mice after 7 days of treatment. Treatments were anti-RW or anti-Jag1/2. All values are expressed as mean ± s.d. and at least n=3 mice per group. Statistical significance was assessed by Mixed-effects analysis with Tukey's multiple comparisons test: $p<0.05$, *; p<0.0001, ****.

F. Serum levels of the indicated cytokines and chemokines in mice treated with anti-RW or anti-Jag1/2 for 2 days. All values are expressed as mean ± s.d. and n=7 mice per group. Statistical analysis was assessed by multiple unpaired *t*-tests.

G. Serum levels of the indicated cytokines and chemokines in mice treated with anti-RW or anti-Jag1/2 for 5 days. All values are expressed as a mean ± s.d. of n=18 mice per group. Statistical analysis was assessed by multiple unpaired *t*-tests.

H. Serum levels of the indicated cytokines and chemokines in mice treated with anti-RW or anti-Jag1/2 for 5 days. All values are expressed as a mean ± s.d. of n=18 mice per group. Statistical analysis was assessed by multiple unpaired *t*-tests.

I. White blood cell counts in mice treated with anti-RW or anti-Jag1/2 at day 7. All values are expressed as mean ± s.d. of at least n=3 mice per group. Statistical significance was assessed using multiple unpaired *t*-tests. NEU, neutrophils; LYM, lymphocytes; MON, monocytes; BAS, basophils.

J. Blood cell counts in mice treated with anti-RW or anti-Jag1/2 at day 7. All values are expressed as mean ± s.d. of at least n=3 mice per group. Statistical significance was assessed using multiple unpaired t-tests. WBC, white blood cells; RBC, red blood cells; HGB, hemoglobin; MCH, mean corpuscular hemoglobin.

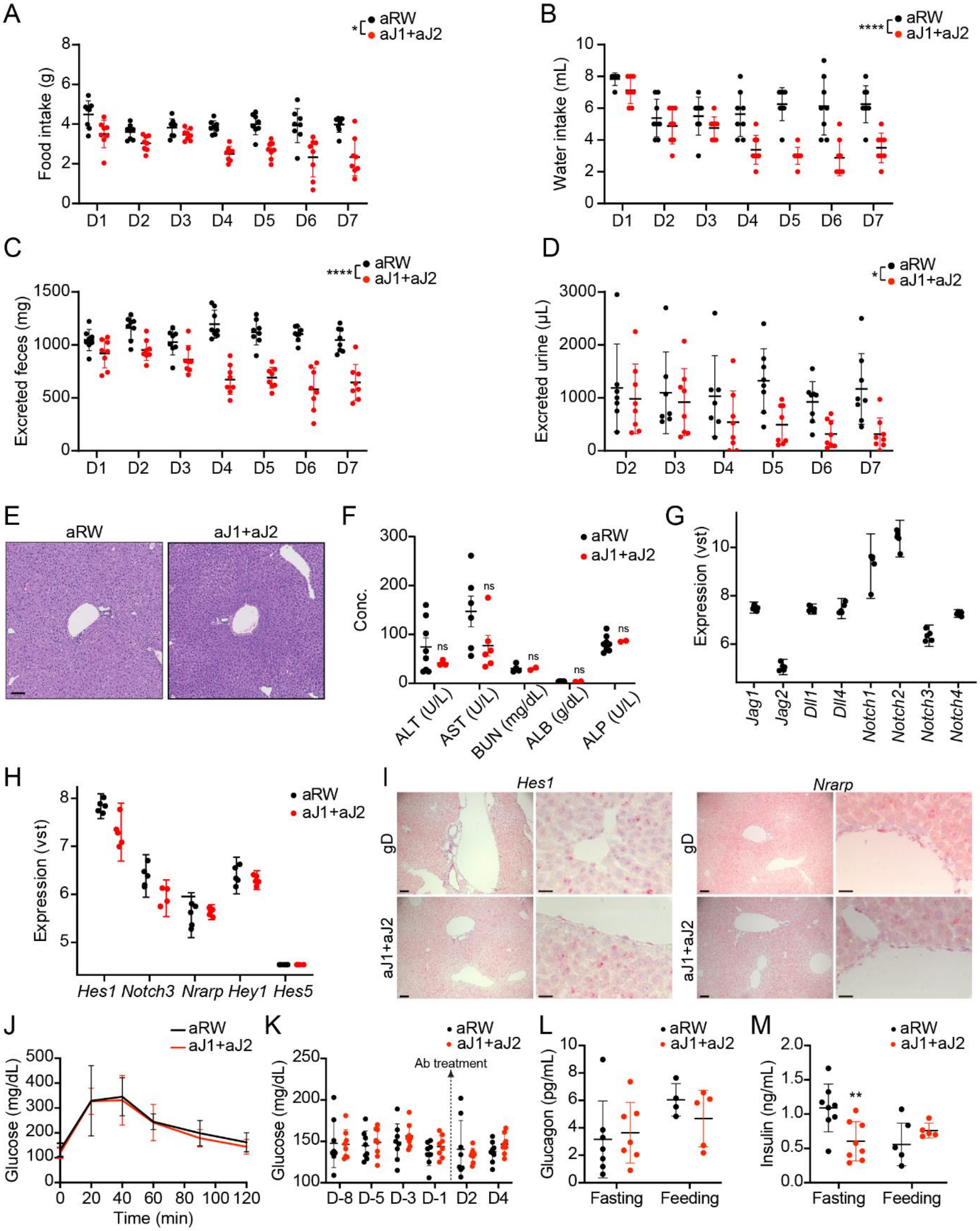


Extended Data Figure 2

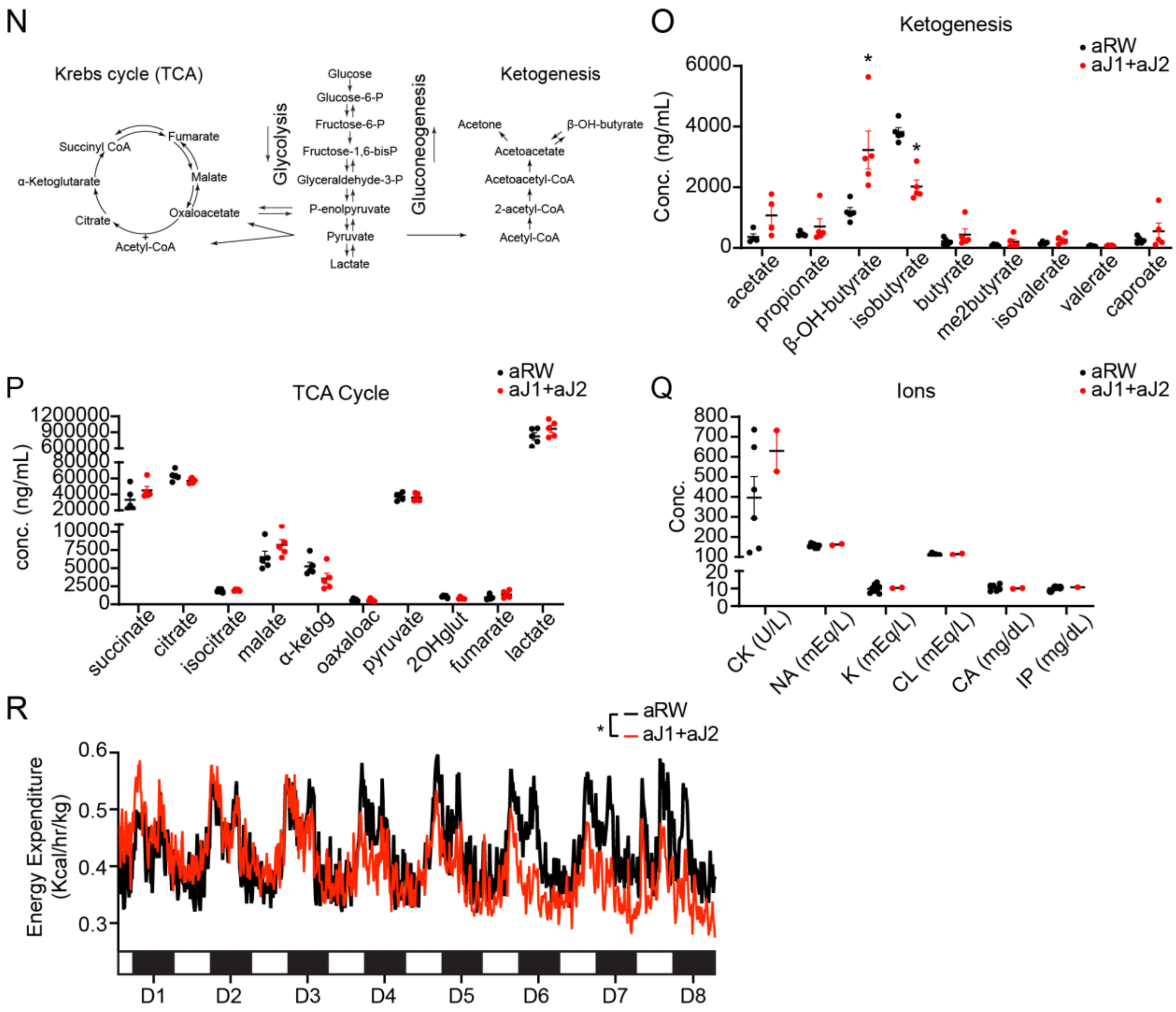


Extended Data Figure 2

**Extended Data Figure 2. Metabolic alterations induced by anti-Jag1/2 treatment.**

A. Daily food intake (in grams, g) of mice treated with anti-RW or anti-Jag1/2 for 7 days. All values are expressed as the mean ± s.d. of at least n=6 mice per group. Statistical significance was assessed using two-way ANOVA, variation tested was treatment over time F=3.876, $p<0.05$, *.
B. Daily water intake (in milliliters, mL) of mice treated with anti-RW or anti-Jag1/2 for 7 days. All values are expressed as the mean ± s.d. of at least n=6 mice per group. Statistical significance was assessed using Mixed-effects analysis, fixed effect tested was treatment over time F=8.578, $p<0.0001$, ****.
C. Daily excretion of feces (in grams, g) of mice treated with anti-RW or anti-Jag1/2 for 7 days. All values are expressed as the mean ± s.d. of n=8 mice per group. Statistical significance was assessed using two-way ANOVA, variation tested was treatment over time F=11.39, $p<0.0001$, ****.
D. Daily excretion of urine (in microliters, µL) of mice treated with anti-RW or anti-Jag1/2 for 7 days. All values are expressed as the mean ± s.d. of at least n=7 mice per group. Statistical significance was assessed using Mixed-effects analysis; the fixed effect tested was treatment over time. F=4.239, $p<0.0001$, ****.
E. Haematoxylin-eosin staining of liver sections of C57BL6 mice under chow diet treated with anti-RW or anti-Jag1/2 for 7 days. Scale bar = 100 µm.
F. Serum levels of the indicated enzymes in paired-fed mice treated with anti-RW or anti-Jag1/2 for 16 hours, 7 days. All values are expressed as the mean ± s.d. of at least n=3 mice per group. Statistical significance was assessed using multiple unpaired *t*-tests. ALB, albumin; ALT, Alanine Transaminase; AST, Aspartate Aminotransferase; ALP, Alkaline Phosphatase; BUN, Blood Urea Nitrogen.
G. Jitter plots showing the variance stabilizing transformation (VST) of RNA-seq counts for Notch pathway components. Liver tissues of mice treated with anti-RW for 7 days. All values are expressed as mean ± s.d. of n=5 mice per group.
H. Jitter plots showing the variance stabilizing transformation (VST) of RNA-seq counts for Notch downstream targets. Liver tissues of paired-fed mice treated with anti-RW or anti-Jag1/2. All values are expressed as mean ± s.d. of n=5 mice per group. Statistical significance was assessed using multiple unpaired *t*-tests.
I. RNAscope of liver of mice treated with anti-RW or anti-Jag1/2 for 7 days. Scale bar = 100 µm (left panel), scale bar = 20 µm (right panel)
J. Measurement of blood glucose levels in a Glucose Tolerance Test (GTT). Treatments were anti-RW or anti-Jag1/2. GTT was performed after 6 hours fasting on day 5 after the antibody treatment. All values are expressed as mean ± s.d. n=8 mice per group. Statistical significance was assessed using two-way ANOVA with Sidak's multiple comparison.
K. Glucose levels in serum of mice 8 days before and 4 days after the treatment. Treatments were anti-RW or anti-Jag1/2. The arrow depicts the start of antibody treatments. All values are

expressed as mean ± s.d. n=8 mice per group. Statistical significance was assessed using two-way ANOVA with Sidak's multiple comparison.

L. Serum glucagon levels in mice treated with anti-RW or anti-Jag1/2 for 7 days. On the left, mice were fasted for 5 hours prior to the measurement. On the right, mice were fed ad libitum. All values are expressed as mean ± s.d. of at least n=4 mice per group. Statistical significance was assessed by multiple unpaired *t*-tests.

M. Serum insulin levels in mice treated with anti-RW or anti-Jag1/2 for 7 days. On the left, mice were fasted for 5 hours prior to the measurement. On the right, mice were fed ad libitum. All values are expressed as mean ± s.d. of at least n=5 mice per group. Statistical significance was assessed by multiple unpaired *t*-tests, $p$adj<0.01, **.

N. Schematic representation of glycolysis and gluconeogenesis, Krebs (TCA) cycle and ketogenesis at the biochemical level.

O. Serum levels of the indicated ketogenesis intermediates in paired-fed mice treated with anti-RW or anti-Jag1/2 for 7 days. All values are expressed as the mean ± s.d. of at least n=6 per group. Statistical significance was assessed using multiple Mann-Whitney tests, $p$adj<0.05, *.

P. Serum levels of the indicated TCA cycle intermediates in paired-fed mice treated with anti-RW or anti-Jag1/2 for 7 days. All values are expressed as the mean ± s.d. of at least n=6 per group. Statistical significance was assessed using multiple Mann-Whitney tests. ɑ-ketog, ɑ-ketoglutarate; 2OHglut, 2-hydroxyglutarate.

Q. Serum levels of the indicated ions in paired-fed mice treated with anti-RW or anti-Jag1/2 for 7 days. All values are expressed as the mean ± s.d. of at least n=2 per group. Statistical significance was assessed using multiple unpaired *t*-tests, $p$adj<0.05, *. CK, creatine kinase; IP, Inorganic Phosphate.

R. Representation of the energy expenditure normalized to body weight of mice treated with anti-RW or anti-Jag1/2 for 8 days. Each cycle represents one day and the bar in the lower part represents the dark cycle (black, nighttime) and light cycle (white, daytime). All values are expressed as mean n=7 mice per group. Statistical analysis was assessed using a mixed effects model, $p<0.05$, source of variation time x genotype, $p<0.001$; ****.

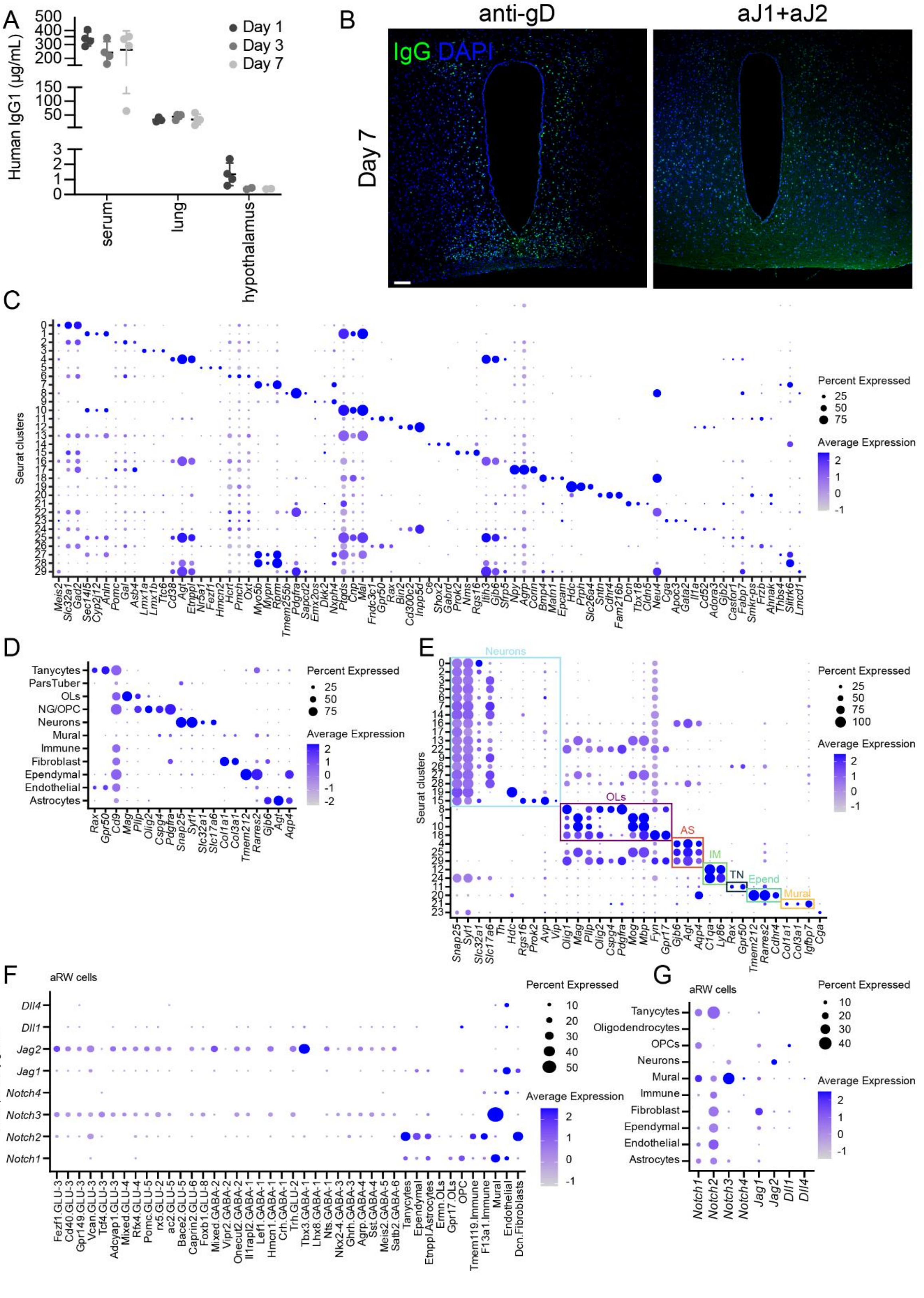


Extended Data Figure 3

**Extended Data Figure 3. Hypothalamic cell types after anti-Jag1/2 treatment**

A. Human IgG1 (anti-Jag1) antibody concentration was measured in serum, lung and hypothalamus at day 1, 3 and 7 post-treatment with anti-Jag1/2 antibodies. n=4 mice per group.
B. Immunostaining against human IgG in the hypothalamus of mice after 7 days after treatment with anti-gD control (human IgG), or anti-Jag1/2 (human IgG). Scale bar=100 µm.
C. Dot plot of the top three markers identified using FindMarkers expressed in each of the 30 clusters defined in the snRNA-seq of the hypothalamus of mice treated for 7 days with anti-RW and anti-Jag1/2 (n=4 mice per group). The size of the dot represents percent of cells, color grade average expression level.
D. Dot plot of canonical cell type markers expressed across annotated cell types. The size of the dot represents percent of cells, color grade average expression level. All values are expressed as the mean of n=4 mice per group.
E. Dot plot of canonical cell type markers expressed across the 30 clusters identified in the snRNA-seq. The size of the dot represents percent of cells, color grade average expression level. All values are expressed as the mean of n=4 mice per group. OL, oligodendrocytes; AS, astrocytes; IM, immune; TN, tanycytes; Expend, ependymocytes.
F. Dot plot of the expression levels of Notch ligands and receptors in the anti-RW samples across the 30 clusters. The size of the dot represents percent of cells, color grade average expression level. All values are expressed as the mean of n=4 mice per group.
G. Dot plot of the expression levels of Notch ligands and receptors in the anti-RW samples across the annotated cell types. The size of the dot represents percent of cells, color grade average expression level. All values are expressed as the mean of n=4 mice per group. OPCs, oligodendrocyte progenitor cells.

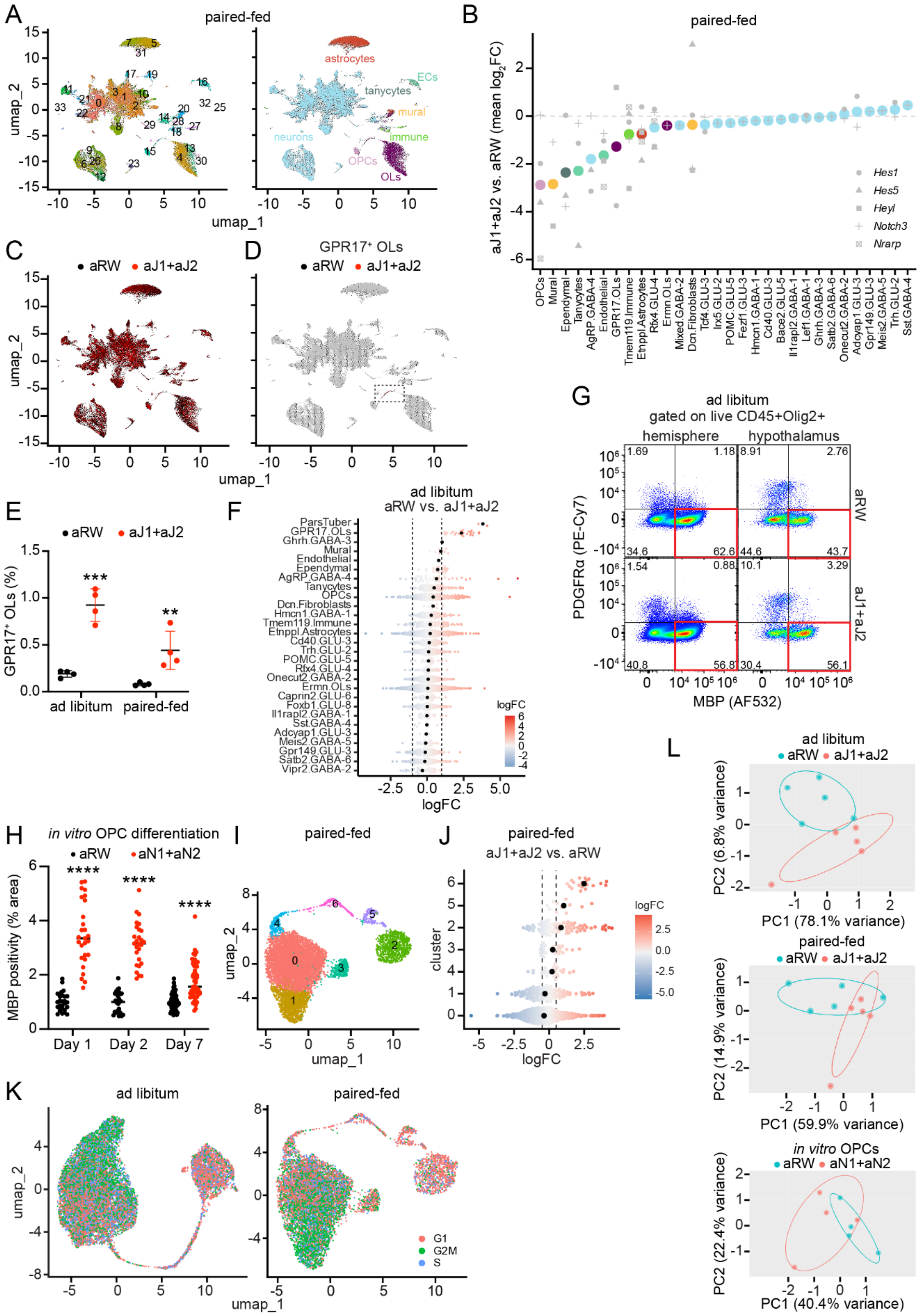


Extended Data Figure 4

**Extended Data Figure 4. Anti-Jag1/2 triggers oligodendrocyte differentiation**

A. UMAP plot and annotated UMAP plot of the snRNA-seq of the hypothalamus of paired-fed mice treated with anti-RW or anti-Jag1/2 for 7 days. Colors highlight the identified clusters on the left UMAP and labels according to HypoMap annotation on the right UMAP. n=4 mice per group. ECs, endothelial cells; OLs, oligodendrocytes; OPCs, oligodendrocyte progenitor cells.

B. Downregulation of Notch target genes by cell type using the C66 labels from HypoMap in the paired-fed dataset. Average downregulation was assessed as a group in anti-Jag1/2-treated compared to anti-RW-treated mice, n=4 mice per group. OLs, oligodendrocytes; OPCs, oligodendrocyte progenitor cells.

C. UMAP plot of snRNA-seq of the hypothalamus of paired-fed mice treated for 7 days with anti-RW (black) and anti-Jag1/2 (red). n=4 mice per group.

D. UMAP highlights the GPR17$^{+}$ oligodendrocyte cluster in red. OLs, oligodendrocytes.

E. Bar plot of the proportions of GPR17$^{+}$ cells in ad libitum and paired-fed datasets. Values are expressed as mean ± s.d. of n=4 mice per group. Statistical significance was assessed using multiple unpaired *t*-test, *p*adj<0.01; **; *p*adj<0.001; ***. OLs, oligodendrocytes.

F. Differential abundance of cells in the hypothalamus of ad libitum fed mice treated with anti-Jag1/2 compared to anti-RW for 7 days, analyzed by MiloR. Number of animals n=4 per group. OLs, oligodendrocytes; OPCs, oligodendrocyte progenitor cells.

G. Flow cytometry plot of oligodendrocyte progenitor cells (PDGFRα+) and mature oligodendrocytes (MBP$^{+}$) in the hypothalamus of ad libitum fed mice treated with anti-RW or anti-Jag1/2 for 7 days, n=5 mice per group.

H. Quantification of the percentage of MBP$^{+}$ area normalized to total Olig2$^{+}$ cells after *in vitro* differentiation of primary OPCs co-cultured with astrocytes, treated with anti-RW or anti-Notch1/2. All values are expressed as mean ± s.d. of at least n=27 images per group. Statistical significance was assessed using unpaired multiple *t*-test, padj<0.0001, **** .

I. UMAP of the oligodendrocyte lineage in the paired-fed datasets. n=4 mice per group.

J. Differential abundance of cells in the oligodendrocyte lineage of paired-fed mice treated with anti-Jag1/2 compared to anti-RW for 7 days, analyzed by MiloR. Number of animals n=4 per group.

K. Cell cycle analysis in the UMAP of the oligodendrocytes lineage in ad libitum and paired-fed datasets. Colors represent the cell cycle phases (G1, G2M, S).

L. Principal Component Analysis (PCA) of metabolomic results in the hypothalamus of ad libitum-fed or paired-fed mice treated with anti-RW or anti-Jag1/2 for 7 days (n=5 mice per group) and PCA of metabolomic analysis of isolated OPCs treated with anti-Notch1/2 for 3 days (n=4 wells per group).

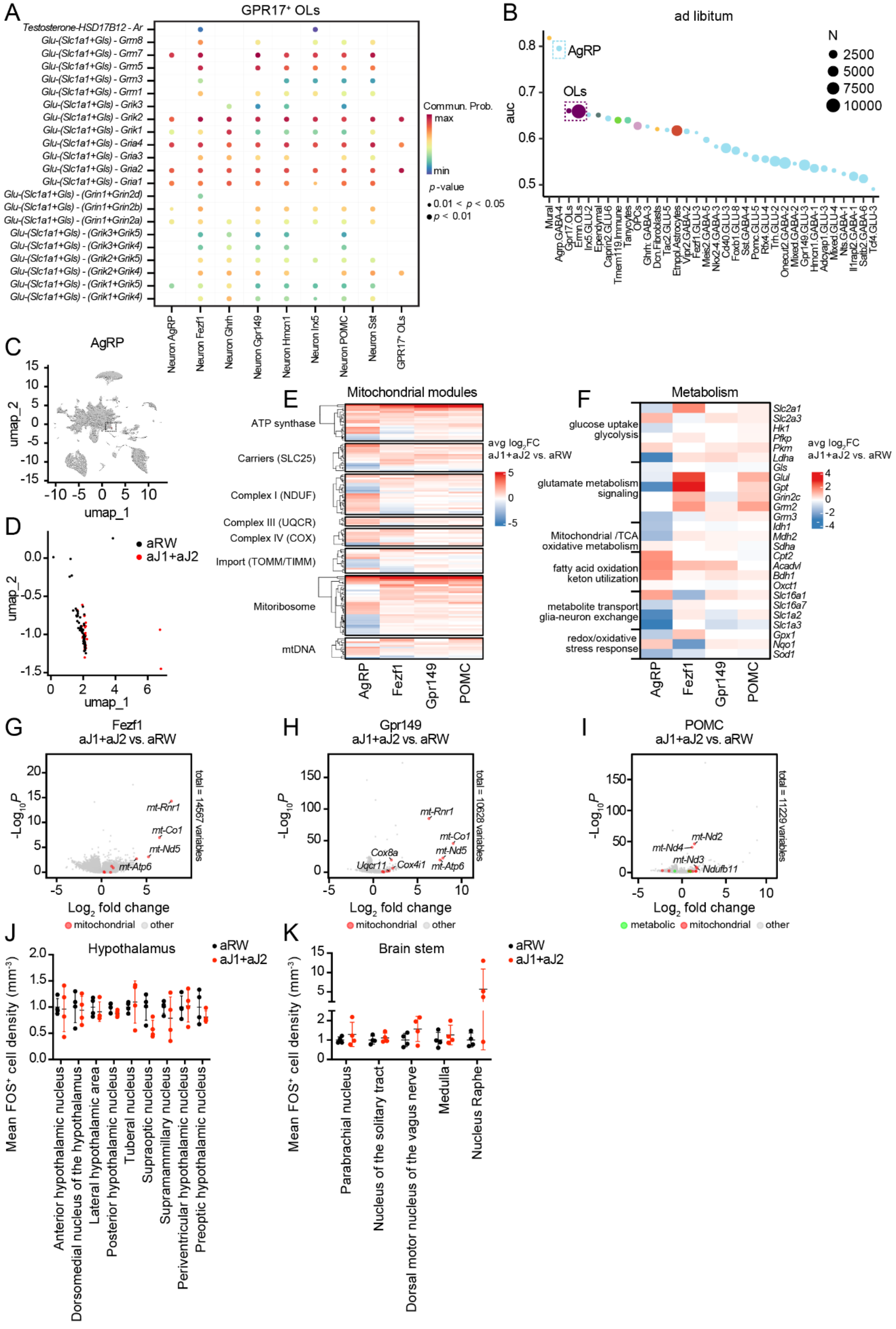


Extended Data Figure 5

**Extended Data Figure 5. Increased communication between GPR17$^+$ oligodendrocytes and neurons results in metabolic changes**

A. Dot plot of ligand-receptor communication pairs (Y axis) in the glutamate pathway identified by CellChat using the non-proteins interactome database. Ligands are expressed in GPR17$^+$ oligodendrocytes and receptors in either neurons or GPR17$^+$ oligodendrocytes (X axis). Circles size represents *p* value and circle color represents communication probability. OLs, oligodendrocytes.
B. Augur analysis in the 30 clusters identified in the snRNA-seq of the hypothalamus of ad libitum mice treated for 7 days with anti-RW or anti-Jag1/2 and annotated according to C66 HypoMap using label transfer. Clusters with cell numbers < 20 were excluded from the analysis. Y axis represents the Area Under the Curve (AUC) of the transcriptomic changes between treatments. Dot size indicates the number of cells per cluster. Color represents annotated cell types. n=4 mice per group. OLs, oligodendrocytes; OPCs, oligodendrocyte progenitor cells.
C. UMAP of snRNA-seq of the hypothalamus of paired-fed mice treated for 7 days with anti-RW or anti-Jag1/2, highlighting the AgRP neurons in red.
D. UMAP magnification AgRP neurons in (C).
E. Heatmap of differentially deregulated mitochondrial genes in anti-Jag1/2- versus anti-RW-treated paired-fed mice in the AgRP, Fezf1, GPR149 and POMC neurons. n=4 mice per group.
F. Heatmap of differentially deregulated genes related to glycolysis, glutamate, TCA/oxidative metabolism, FAO/ketones, metabolite transport and redox in anti-Jag1/2- versus anti-RW-treated paired-fed mice in the AgRP, Fezf, GPR149 and POMC neurons. n=4 mice per group.
G. Volcano plot of the differentially expressed genes in anti-Jag1/2- compared to anti-RW-treated mice in Fezf1 neurons in the hypothalamus of paired-fed mice treated for 7 days, n=4 mice per group. Mitochondrial genes are highlighted in red.
H. Volcano plot of the differentially expressed genes in anti-Jag1/2- compared to anti-RW-treated mice in GPR149$^+$ neurons in the hypothalamus of paired-fed mice treated for 7 days, n=4 mice per group. Mitochondrial genes are highlighted in red.
I. Volcano plot of the differentially expressed genes in anti-Jag1/2- compared to anti-RW-treated mice in POMC neurons in the hypothalamus of paired-fed mice treated for 7 days, n=4 mice per group. Genes related to metabolism are highlighted in green and mitochondrial genes in red.
J. Quantification of TdTomato cell density in the hypothalamic nuclei of Fos-Trap (cFos-CreERT2; LSL-tdTomato) mice treated with anti-RW or anti-Jag1/2 for 4 days. Number of animals n=4 per group. Statistical significance was assessed using multiple Mann-Whitney tests.
K. Quantification of TdTomato intensity in the melanocortin output nuclei of the brainstem in Fos-Trap (cFos-CreERT2; LSL-tdTomato) mice treated with anti-RW or anti-Jag1/2 for 4 days, n=4 mice per group. Statistical significance was assessed using multiple Mann-Whitney tests.

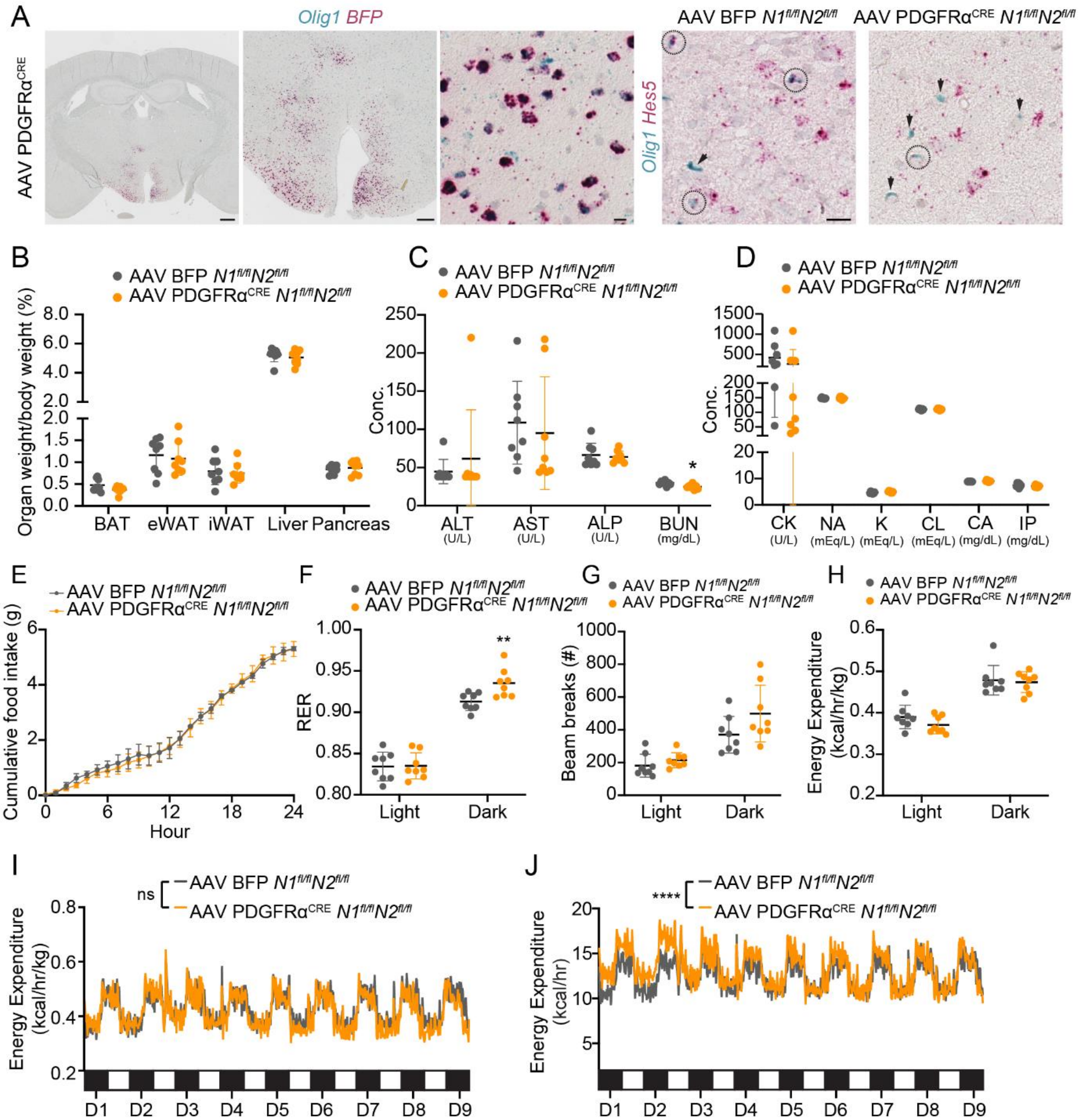


Extended Data Figure 6

**Extended Data Figure 6. OPC-specific deletion of *Notch*1/2 in the hypothalamus recapitulates metabolic effects of anti-Jag1/2 treatment**

A. RNAscope of Blue Fluorescent Protein (*BFP*) in red and *Olig1* in blue in the hypothalamus of AAV.PdgfrαCre$^{+}$; *Notch1*$^{fl/fl}$*; Notch2*$^{fl/fl}$ mice. RNAscope of *Hes5* in red and *Olig1* in blue in the hypothalamus of AAV.Cre$^{-}$; *Notch1*$^{fl/fl}$*; Notch2*$^{fl/fl}$ mice or AAV.PdgfrαCre$^{+}$; *Notch1*$^{fl/fl}$; *Notch2*$^{fl/fl}$ mice. Left panel scale bar = 200 µm, middle panel scale bar = 10 µm, right panels scale bar = 20 µm. Arrows mark *Olig1*$^{+}$ *Hes5*$^{-}$ cells and circles mark *Olig1*$^{+}$*Hes5*$^{+}$ cells.

B. Percentage of wet tissue weight compared to initial body weight of mice in (Fig 6A). All values are expressed as mean ± s.d. of at least n=8 mice per group. Statistical significance was assessed using multiple unpaired *t*-tests. BAT, brown adipose tissue; eWAT, epididymal white adipose tissue; iWAT, inguinal white adipose tissue.

C. Serum levels of the indicated enzymes in mice from (Fig 6A) assessed at day 59. All values are expressed as the mean ± s.d. of n=8 mice. Statistical significance was assessed using multiple unpaired *t*-tests. ALT, Alanine Transaminase; AST, Aspartate Aminotransferase; ALP, Alkaline Phosphatase; BUN, Blood Urea Nitrogen.

D. Serum levels of the indicated ions in mice from (Fig 6A) assessed at day 59. All values are expressed as the mean ± s.d. of n=8 per group. Statistical significance was assessed using multiple unpaired *t*-tests. CK, creatine kinase; IP, Inorganic Phosphate.

E. Daily food intake in mice from (A), measured during 24h at day 49 after stereotaxic injection. All values are expressed as the mean ± s.d. of at least n=8 mice per group. Statistical significance was assessed by Mixed effects analysis.

F. Representation of the respiratory exchange ratio (RER) of mice from (Fig 6A) during light and dark time measured 47-56 days after stereotaxic injection. The graph represents the average of total measurements in light and dark cycles. All values are expressed as an average of n=8 mice per group. Statistical significance was assessed using multiple unpaired *t*-test, $p$adj<0.01, **.

G. Representation of the mobility (assessed by number of beam breaks) of mice from (Fig 6A) during days 47-56 after stereotaxic injection. The graph represents the average of total measurements in light and dark cycles. All values are expressed as an average of n=8 mice per group. Statistical significance was assessed using multiple unpaired *t*-tests.

H. Representation of the absolute energy expenditure of mice from (Fig 6A) during days 47-56 after stereotaxic injection. The graph represents the average of total measurements in light and dark cycles. All values are expressed as the mean of n=8 mice per group. Statistical significance was assessed using multiple unpaired *t*-tests.

I. Representation of the energy expenditure relative to body weight of mice from (Fig 6A) during days 47-56 after stereotaxic injection. Each cycle represents one day and the bar in the lower part represents the dark cycle (black, nighttime) and light cycle (white, daytime). All values are expressed as the mean of n=8 mice per group. Statistical analysis was assessed using a mixed effects model.

J. Representation of the absolute energy expenditure of mice from (Fig 6A) during days 47-56 after stereotaxic injection. Each cycle represents one day and the bar in the lower part represents the dark cycle (black, nighttime) and light cycle (white, daytime). All values are expressed as the mean of n=8 mice per group. Statistical analysis was assessed using a two-way ANOVA, source of variation time x genotype, $p<0.001$; ****.
